# Disruption of ubiquitin mediated proteolysis is a widespread mechanism of tumorigenesis

**DOI:** 10.1101/507764

**Authors:** Francisco Martínez-Jiménez, Ferran Muiños, Erika Lopez-Arribillaga, Nuria Lopez-Bigas, Abel Gonzalez-Perez

## Abstract

E3 ligases and degrons --the sequences they recognize in target proteins-- are key parts of the ubiquitin-mediated proteolysis system. There are several examples of alterations of these two components of the system that play a role in cancer. Here, we uncovered the landscape of the contribution of such alterations to tumorigenesis across cancer types. We first systematically identified novel instances of degrons across the human proteome using a random forest classifier, and validated them exploiting somatic mutations across more than 7,000 tumors. We detected signals of positive selection across these novel degrons and revealed new instances involved in cancer development. Overall, we estimated that at least one in seven driver mutations across primary tumors affect either degrons or E3 ligases.

## Introduction

The targeted degradation of proteins *via* the ubiquitin-mediated proteolysis system (UPS), constitutes a key step in the maintenance of protein levels for intracellular homeostasis (Guharoy et al., 2016; Liu et al., 2016; Mészáros et al., 2017). The UPS is involved in both quality control – eliminating misfolded proteins– and spatio-temporal control of the level of proteins that regulate crucial processes, such as the advance of cell cycle (Bassermann et al., 2014), the circadian clock (Yoo et al., 2013) or DNA repair (Al-Hakim et al., 2010; Arlow et al., 2013; Gillette et al., 2006). The recognition of specific targets for degradation is achieved through the interaction of E3-ubiquitin ligases, or E3s, for short (of which the human genome contains around 600) and short sequences within the target called degrons. While N-terminal and C-terminal degrons are short sequences (often only one or two amino acids), which are exposed by the action of exopeptidases, internal degrons are sequences spanning 3 to 10 amino acids, frequently flanked by phosphorylation sites. The focus of this study is internal degrons, which we will from here on out refer to as degrons for simplicity. Since several related sequences may be recognized by an E3, degrons –much like transcription factors binding sites– are degenerate. We thus use the term motif to denote the representation (as a regular expression) of the collection of sequences that represent a degron. We refer to each of these sequences as a degron instance.

Upon binding of a degron instance by the cognate E3, a second enzyme (E2, ubiquitin conjugating) transfers a ubiquitin –previously loaded onto it by the E1 activating enzyme– to a lysine of the target protein. This catalysis occurs within a complex formed by the specific E3, one E2 (out of the 41 encoded in the human genome), and the target protein (Stewart et al., 2016). Such complexes may be completed by scaffolding, adaptor and/or substrate receptor proteins. Substrate lysines can be tagged by a single ubiquitin molecule (monoubiquitination) or by several concatenated ubiquitins (polyubiquitination). Typically polyubiquitin chains linked through residues K-48 or K-11 of the ubiquitin polypeptide work as the signal for proteasomal degradation, although this can also be triggered following monoubiquitination (Braten et al., 2016). The system also comprises erasers (deubiquitinases) that counteract the activity of the E2-E3 complexes (Komander et al., 2009).

Examples of dysregulation of the UPS that result in changes of the stability of certain proteins have long been linked to human diseases (Vu and Sakamoto, 2000). There are, for example, well-known degron-affecting mutations in oncoproteins, such as CTNNB1 and NFE2L2 (Mészáros et al., 2017), and loss-of-function alterations of E3 ligases, such as FBXW7, SPOP and APC that appear recurrently across certain types of cancer (Ge et al., 2018). Here, we asked how pervasive across tumorigenesis the alteration of UPS mechanisms that lead to abnormal stabilization of oncoproteins are. A thorough landscape of the involvement of UPS dysregulation in cancer has been not possible to date due to the paucity in the identification of E3-target links. Three decades of laborious biochemical studies since the discovery of the UPS in the 1980s (Ciechanover et al., 1980; Hershko et al., 1980) have yielded the identification of 150 annotated instances of 29 degron motifs across 98 proteins (Dinkel et al., 2016; Mészáros et al., 2017). Since this represents less than 5% of all E3 ligases, the number of annotated degrons is likely a small fraction of all degrons in the human proteome.

To study the involvement of abnormal oncoprotein stabilization via UPS alteration in tumorigenesis, we first needed to identify new degron-mediated E3-target links. Because the sequence motifs of known degrons are short and degenerate, simple pattern matching methods are expected to produce a long list of potential matches enriched for false positive hits. To circumvent this problem, we designed a random forest classifier that identifies new instances of degrons based on their biochemical features. Experimental validation of these novel degron instances would imply carrying out site-directed mutagenesis of each of them and direct measures of protein stability change. To carry out such validation here, we exploited somatic coding mutations in tumors as a “natural experiment” of mutagenesis. By controlling for the mRNA influence on protein concentration, we were able to identify mutations that significantly increase the stability of proteins, and further used this highly stabilizing mutations to validate novel –and find *de novo–* E3-target links. We then used these links to identify degrons under positive selection in tumorigenesis and to identify the contribution of E3 and adaptors to cancer. Overall, we estimated that alterations of UPS elements (degron instances and E3s) contribute one in seven driver mutations across primary tumors.

## Results

### New instances of known degrons identified with a machine learning approach

Experimentally identified degron instances share several common biochemical features (Guharoy et al., 2016). We thus reasoned that these properties could be learned by means of an appropriate statistical method to identify new instances across the human proteome. To that end, we systematically explored how much a set of biochemical features –inspired by previous studies concerning smaller sets of degrons (Guharoy et al., 2016; Mészáros et al., 2017)– distinguish a set of 150 known degron instances (Table S1) from sequences of the same length randomly drawn from the human proteome. We found that the properties that best distinguished degrons from other sequences were: disorder, solvent accessibility, rigidity, secondary structure, conservation, anchoring potential, location within structured domains, presence of flanking phosphorylation sites and ubiquitination sites (Fig. 1a,b and Fig. S1a). The average degron is enriched for regions with high disorder and solvent accessibility, flanked by phosphorylation sites and ubiquitinated lysines more frequently than other sequences. It appears in regions with coiled structure more often than other sequences, it is more likely to gain stability upon protein interaction (anchor), and its sequence is more conserved compared to its flanks. On the other hand, it is comparatively depleted of stretches of alpha-helix or beta-strand, it is seldom located within globular domains, and its sequence does not provide high rigidity to the peptide backbone. The importance of particular features differs between types of degrons (Fig. 1b). The sequences of degron instances, in general, are significantly enriched in aspartic acid, proline and serine residues, and they are significantly depleted of cysteines, lysines, leucines and alanines (Fig. S1b).

**Figure 1.**
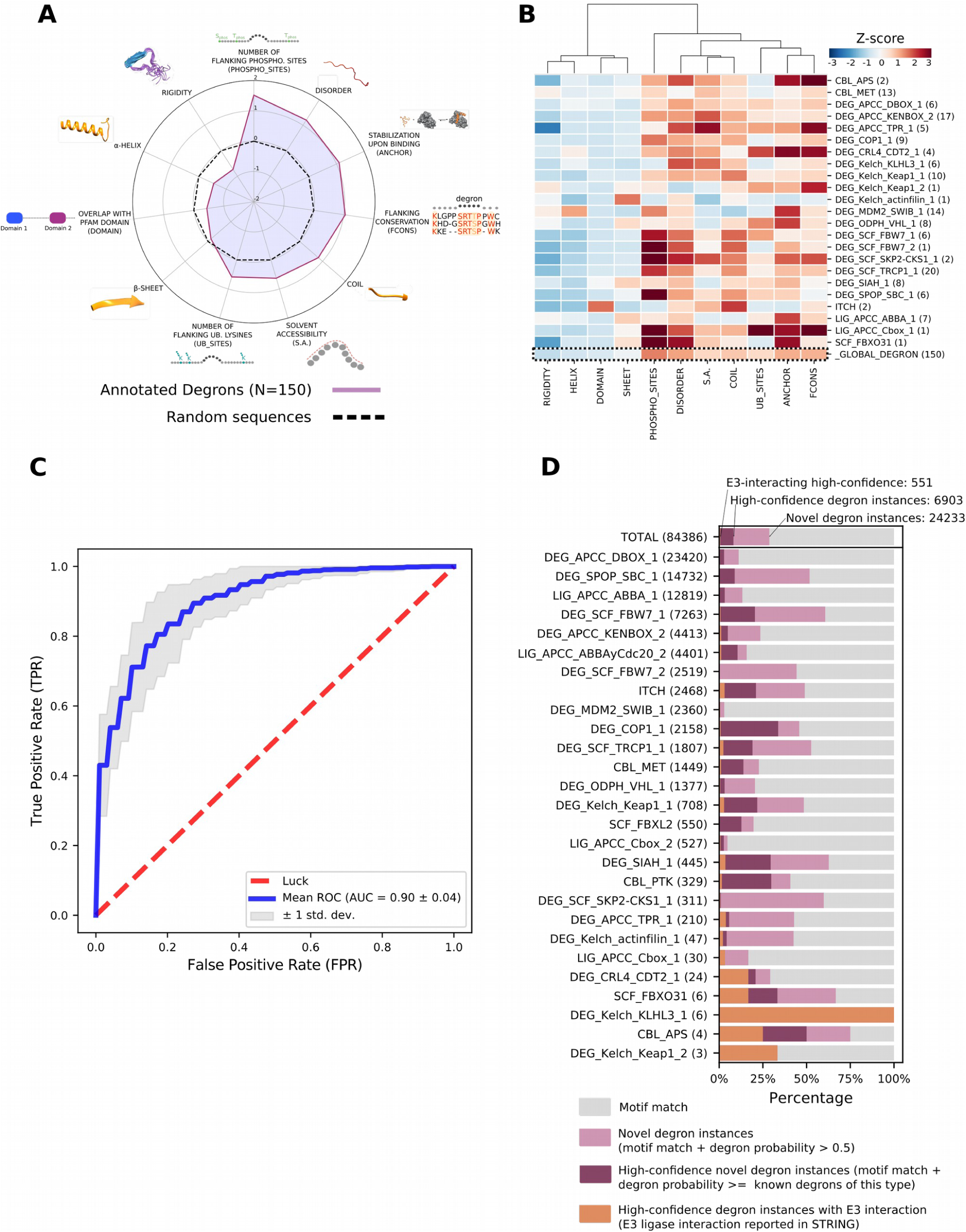
Identification of new instances of known degrons. (a and b) Biochemical features used to differentiate annotated instances of degrons from random protein sequences of the same length. The Z-score of the values of each feature are represented globally in a radial plot (a) and degron-wise in a heatmap (b). (c) Stratified five-fold cross-validation ROC curve of a random forest classifier trained on annotated instances of degrons and random protein sequences. (d) Motif matches (number at the left of the bar) identified for each degron motif. The bars are segmented to represent, for the matches of each degron, the proportion of i) novel degron instances (identified by the classifier in the human proteome with a probability above 0.5; light purple segment), ii) high-confidence novel degron instances (above threshold selected for each degron; dark purple segment), and iii) E3-interacting high-confidence degron instances (high-confidence instances in proteins with evidence of interaction with the cognate E3; orange segment).

We then trained a random forest classifier that predicts how much an amino acid sequence resembles the properties of annotated degrons. The classifier was trained using the set of known degron instances as positive set and the same number of random sequences of comparable lengths obtained from all annotated protein isoforms of the human proteome (Bateman et al., 2017) as negative set, with the 11 biochemical features analyzed above. Ten rounds of 5-fold cross-validation yielded an average area under the receiver operating characteristic (ROC) curve of 0.9 (Fig. 1c). We obtained a similar result with a random forest trained on the same positive set and an alternative negative set composed of random sequences drawn from the same proteins (Fig. S1c,d). The disorder, the anchoring capability, and the sequence conservation of the flanking regions of degrons were the structural features deemed most informative by the classifier in both settings (Fig. S1e,f).

We then scanned the entire human proteome for known degron motifs, i.e., regular expressions that combine all known instances of each known degron. We identified 84386 matches of known degron motifs (motif match in Fig. 1d). Numbers in the left of the barplot in Figure 1d correspond to the matches identified for each degron motif. The number of matches of degrons per protein correlated with the length of the proteins (Fig. S1g), supporting the notion that they contain many false positive instances. The matches of each degron motif may be ranked following the probability predicted by the random forest classifier (degron probability). In this ranking, the higher the degron probability, the more the match resembles the profile of biochemical features of the degron. A generic threshold of degron probability greater than 0.5 is attained by 24233 matches in different protein isoforms (novel degron instances in Fig. 1d). (Only 18239 out of these novel degron instances, can be mapped to Ensembl transcripts; See Methods.) We reasoned that specific thresholds could be set to identify high-confidence novel instances of each degron, based on the distribution of probabilities of all its known instances (Fig. S2). Overall, this subset of high-confidence degron instances comprises 6903 (8.1%) matches (Fig. 1d). For example, we identified 14732 matches of the SPOP degron (SPOP_SBC_1) motif due to its lowly specific regular expression ([AVP].[ST][ST][ST]), including 7618 (51%) novel degron instances (degron probability of degron greater than 0.5). However, only 1310 (8.89%) matches passed the motif-specific threshold (i.e., DEG_SPOP_SBC_1 = 0.84 of degron probability) and were thus considered novel DEG_SPOP_SBC_1 degron instances (See Table S1 for full description and details of degrons analyzed).

Since E3s and the proteins they recognize through specific degrons must come into physical contact, we reasoned we could further refine the set of high-confidence degron instances using protein-protein interaction information (Franceschini et al., 2013). We labeled as “E3-interacting high-confidence degron instances” all matches with probability above threshold that occur in proteins known or predicted to interact with their cognate E3s (Fig. 1d). Following the example of the SPOP_SBC_1 motif, this filtering step further reduced the matches to 23 (0.15%). Overall our approach identified 551 E3-interacting high-confidence degron instances (0.65% of the starting 84386 matches), 416 of which are not part of the starting set of annotated instances. These 551 interacting matches occur in 280 different genes associated to 426 different protein isoforms. Interestingly, the number of degrons per protein shows weaker correlation to the length of the proteins containing them than that of all matches (Fig. S1g). All motif matches annotated with all the information described above are presented in Table S2.

In summary, using a random forest classifier we identified novel degron instances of 29 known motifs, the functionality of which we next sought to experimentally validate.

### Validating the functionality of computationally identified degrons

To experimentally validate novel degron instances identified in the previous section, one could introduce mutations altering their sequence within a cell system and compare the stability of the mutant proteins to that of their wild-type counterparts. The underlying rationale is that changes to a degron sequence may affect the efficiency of the interaction with its cognate E3 and, ultimately, change the stability of the proteins carrying it. These sequence changes could be triggered by non-synonymous mutations or in-frame indels.

Instead of using this experimental setting to elucidate the effect of degron sequence changes on protein stability, we employed human tumors as natural mutagenic experiments. To that end, we required that the abundance of the proteins harboring novel degron instances (and RNA level of their transcripts) had been determined in tumors with and without mutations in said degrons. The Cancer Genome Atlas (TCGA) project has sequenced the exome and transcriptome –and measured the abundance of 209 proteins, through reverse phase protein assays, or RPPAs (Li et al., 2013)– of 7663 primary tumors (Ellrott et al., 2018). Similar measurements have been taken within the Cancer Cell Lines Encyclopedia (CCLE) project for 173 proteins across 889 cancer cell lines (Barretina et al., 2012). Using these data we controlled for the effect of mRNA abundance on protein levels to accurately evaluate the impact of individual mutations on protein abundance. (For details, see Methods and Table S3.)

For each protein we related the protein level to the mRNA level by means of a robust regression model built with wild-type samples (as the examples in Fig. 2a-c). Hence, from the mRNA level observed in a sample, an expected protein level may be obtained. To avoid artifacts, the regression computed for each gene filters out samples with high-level amplifications (Gonçalves et al., 2017), non-synonymous mutations and alterations in upstream annotated E3 ligases. We refer to the distance between the observed protein level and its expected value as “raw residual” (Fig. 2a). To make these raw residuals comparable across proteins --by correcting for inherent experimental and biological variability-- we rescaled them to account for the standard deviation of the protein and mRNA levels observed for all wild-type instances of the protein (Supplemental Note). The rescaled residual of a mutant protein can be regarded as the difference between its observed abundance and its expected value given the level of its mRNA. In other words, it represents the stability change related to the mutation (Fig. 2d-f). The comparison of the distribution of the values of stability change of the wild-type and mutant forms of a protein may thus reveal if its abundance increases (TP53 in colorectal tumors, Fig. 2b,e), decreases (CHEK2 in uterine adenocarcinomas, Fig. 2c,f), or remains fundamentally unchanged (CCNE1 in uterine adenocarcinomas, Fig. 2a,d) upon non-synonymous mutations.

**Figure 2.**
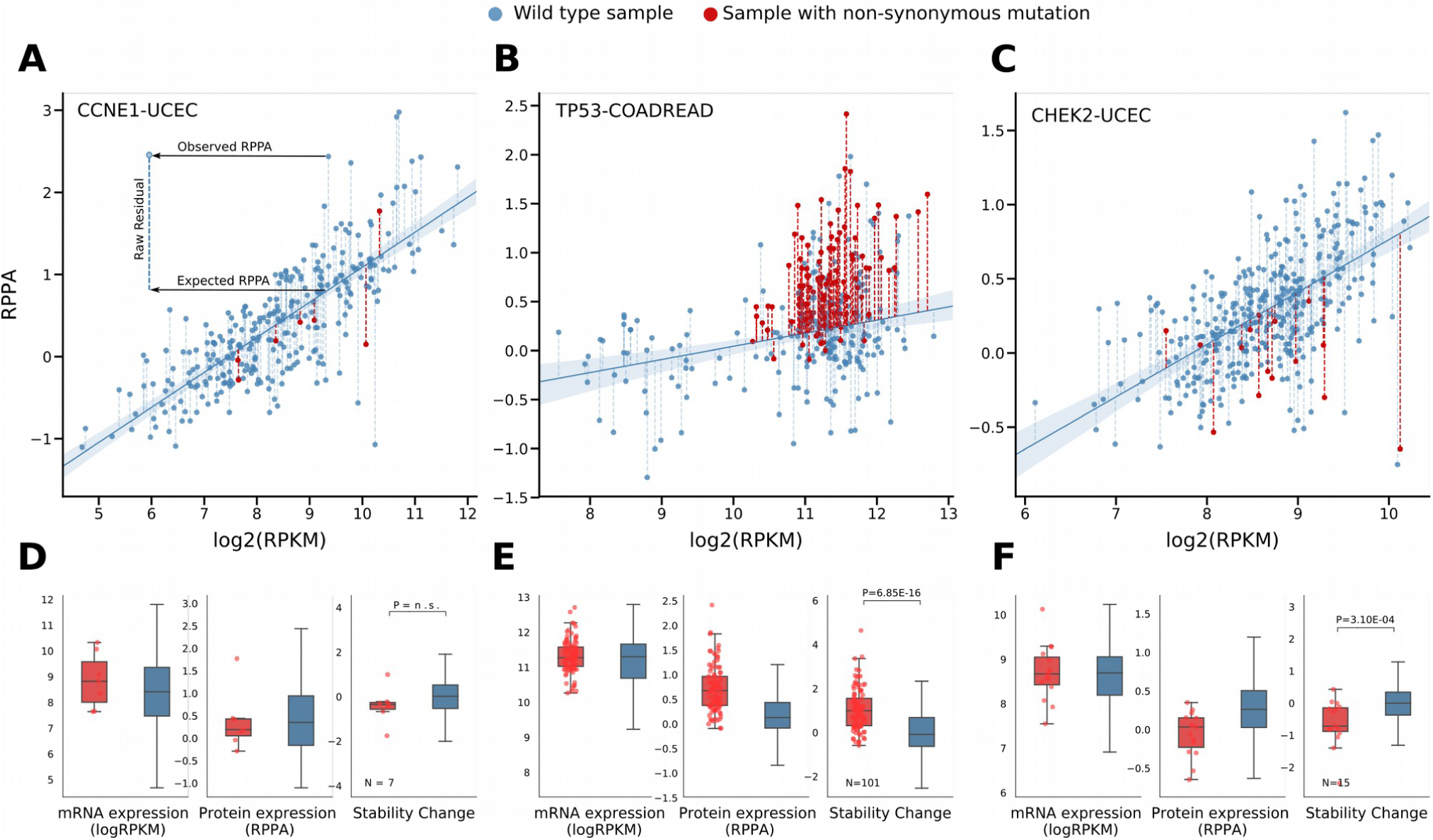
The effect of non-synonymous mutations on the stability of proteins. (a to c) The raw residual of the protein level with respect to its expected value is computed as the distance to a weighted regression between the level of mRNA of the transcript and the protein level of its polypeptide product. The raw residual of mutated instances (red dots) of the protein is a measure of the effect of the mutation on protein stability. (d to f) Comparison (using the Mann-Whitney test, as in subsequent plots) of --from left to right-- the mRNA level, the protein level and the normalized residuals (stability change) of the mutant and wild-type forms of the three proteins represented in panels a-c. The value represented by the y-axis of each plot is indicated below the graph.

To assess the usefulness of this metric of stability change, we analyzed the mutations affecting known degrons of KEAP1 in (Shibata et al., 2008) NFE2L2 and βTRCP inTRCP in (Liu et al., 1999) CTNNB1. There are 40 (26 unique) non-synonymous mutations or in-frame indels in the protein sequence of NFE2L2 across primary tumors, 38 (24 unique) of which affect one of the two Kelch-KEAP1 degrons or their flanking residues (needle-plot in Fig. 3a). The latter probably interfere with the recognition of the degron by KEAP1, probably hindering the ubiquitination reaction. For example, the mutation D29H, abrogates two hydrogen bonds between the N-terminal degron and its binding pocket in KEAP1 (Fig. 3a, top panel). The stability of NFE2L2 is significantly higher in the 38 tumors carrying non-synonymous mutations or in-frame indels (pink dots and triangles, respectively and pink boxplot; Fig. 3b) that affect the two Kelch-KEAP1 degrons and their flanking positions than in the 473 carrying the wild-type protein (blue boxplot; Fig. 3b).

**Figure 3.**
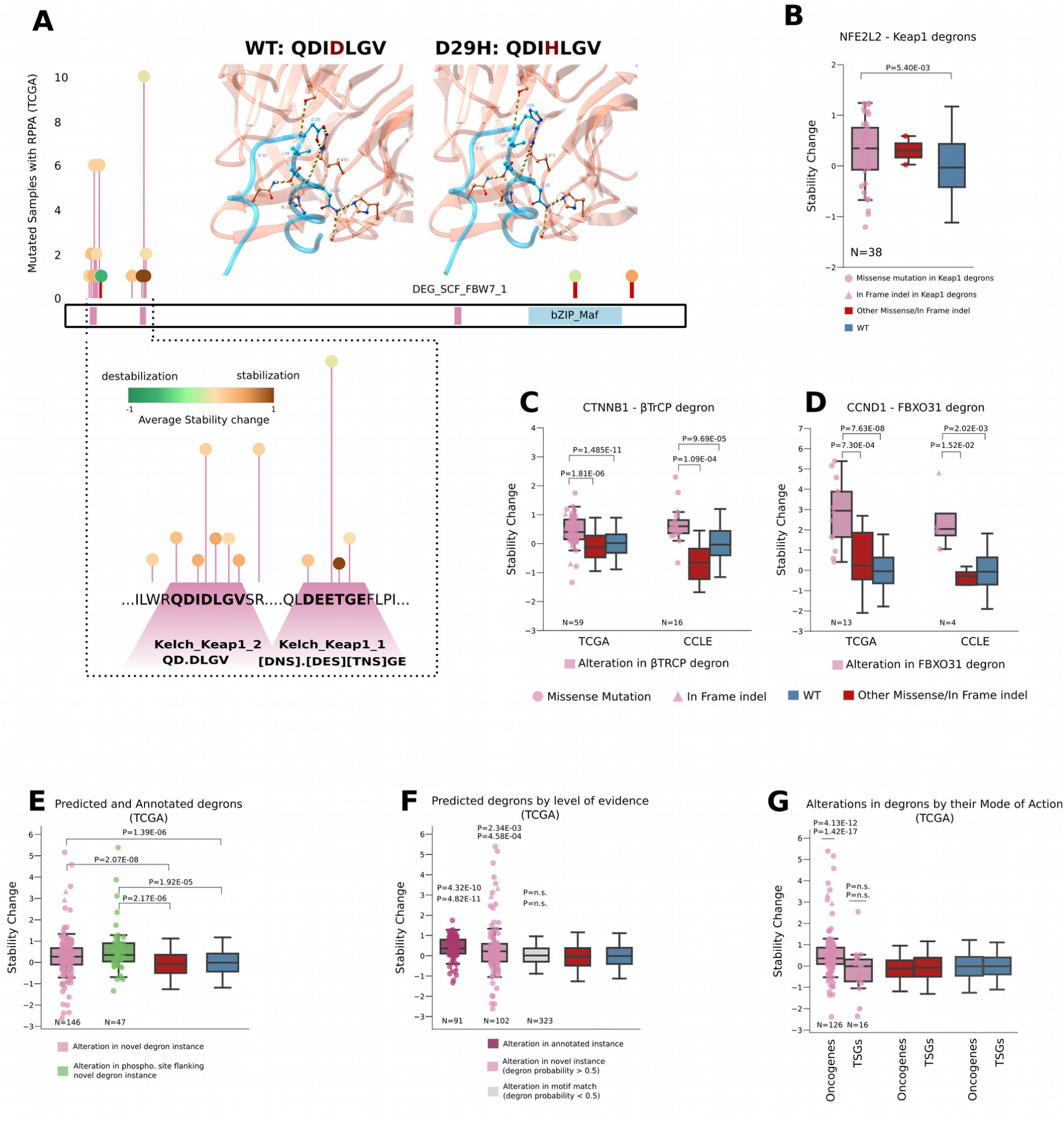
Mutations affecting degrons increase the stability of proteins. (a) Primary tumor mutations of NFE2L2 cluster in its two Kelch_Keap1 degrons, the majority of which result in an increase of the protein stability. Certain positions of the two degron motifs are frequently mutated (zoom-in panel below the mutation needle-plot). Top panels: one of these recurrent mutations (a change of a histidine for an aspartic acid) abrogates two hydrogen bonds (yellow dashed lines) in the interface of interaction between KEAP1 and the degron of NFE2L2. Overall, mutations in Kelch_Keap1 (b), βTRCP (c), and predicted FBXO31 (d) degrons significantlyTRCP (c), and predicted FBXO31 (d) degrons significantly increase the stability of NFE2L2 (b), CTNNB1 (c) and CCND1 (d), respectively. Chimera (Pettersen et al., 2004) software was used to visualize the NFE2L2 KEAP1 interaction (PDB: 3WN7). (e, f) Collectively, mutations in annotated and novel instances (with classifier probability above 0.5) of known degrons or in their flaking phosphosites significantly increase the stability of their proteins (e) The increase of stability is elicited by mutations of both annotated and novel instances, while mutations in sequence matches with lower probability (below 0.5) do not result in protein stability increase (f). (g) Mutations in novel instances identified in known oncogenes significantly increase their stability in comparison with other mutations in the same proteins or their wild-type form. Mutations in novel instances identified in known tumor suppressor genes (TSGs) do not impact their stability in comparison with other mutations in the same proteins or their wild-type form. In all comparisons, TP53 mutations, known to trigger a stabilization of its protein product (Lukashchuk and Vousden, 2007; Wawrzynow et al., 2018) were filtered out.

Analogously, 59 primary tumors and 16 cancer cell lines harbor 22 unique non-synonymous mutations and 11 in-frame indels affecting the CTNNB1 βTRCP inTRCP degron (Fig. S3a). The most frequently affected residue is the serine 37, which is often mutated to a cysteine, a change that results in the abrogation of three hydrogen bonds between the CTNNB1 degron and its binding pocket within βTRCP inTRCP (Fig. S3b). Again, two flanking positions containing a threonine (T41) and a serine (S45), susceptible to phosphorylation also bear non-synonymous mutations that change the abundance of the protein. The stability of CTNNB1 in tumors with non-synonymous mutations or in-frame indels affecting the βTRCP inTRCP degron is significantly higher than in those with the wild-type counterpart, but also in those bearing other non-synonymous mutations in CTNNB1, both across primary tumors and cancer cell lines (red boxplots; Fig. 3c).

Non-synonymous mutations and in-frame indels affecting a novel FBXO31 degron instance in CCND1 (Table S2) –experimental evidences of which exists (Santra et al., 2009)– show the same stabilizing effect (Fig. 3d). CCND1 in tumors carrying non-synonymous mutations or in-frame indels in the novel FBXO31 degron instance (13) (pink boxplots; Fig 3d) shows significantly higher stability than that measured in other CCND1 mutant and wild-type tumors. Four non-synonymous mutations and in-frame indels observed in this degron instance across cancer cell lines also result in stabilization of CCND1.

We then tested whether, globally, the nonsynonymous substitutions and in-frame indels affecting novel degron instances resulted in higher stabilization of the mutant proteins than other mutations. We found that, across primary tumors, 146 mutant proteins carrying these two types of degron-affecting mutations exhibit significantly higher stability than other mutant proteins and wild-type proteins (pink boxplot; Fig. 3e; Methods). Proteins bearing non-synonymous mutations or in-frame indels affecting phosphorylation residues close to novel degron instances (with probability greater than 0.5) are also significantly more stable than other mutant proteins and all wild-type proteins (green boxplot; Fig. 3e). This significant increase of stability is registered for both, annotated and novel instances of degrons (Figs. 3f, S3c-h). Furthermore, the analysis of mass-spectrometry (MS) measured protein abundance across 105 breast and 175 ovary primary tumors revealed that 13 mutations affecting degron matches resulted in an average stabilization of the proteins higher than that registered for other non-synonymous mutations (Fig. S3i). Nevertheless, due to the low numbers of mutated novel degron instances, the comparison of stability increase was not statistically significant. Taken together, these results support the notion that the machine learning-based approach described above produces matches enriched for true instances of annotated degrons.

We also asked whether mutations affecting degron instances of oncogenes and tumor suppressors exhibited different stabilization patterns. We found that while mutations affecting annotated and novel degron instances of oncogenes result in significantly higher stabilization than other mutations in the same proteins or their wild-type form, mutations in known and newly identified degrons of tumor suppressors do not result in any increase or decrease of their stability (Fig. 3g). Most of the global increase of protein stability resulting from degron mutations in tumors can therefore be explained by known oncogenes.

In summary, exploiting naturally occurring mutations in tumors, we provide *in vivo* experimental validation of the functionality of novel degron instances.

### Highly stabilizing mutations reveal potential *de novo* degrons

Degron motifs have been identified for less than 5% of the 600 E3s estimated to be encoded in the human genome; therefore, many degrons remain to be discovered. We reasoned that highly stabilizing mutations (represented as dots in the top quartile of stability change values in panel 1 of Fig. 4a) that do not overlap degron matches (red dots) could affect degrons with still unknown motif. To discover some of these *de novo* degrons, we searched for proteins with regions of high degron probability bearing recurrent highly stabilizing mutations. Briefly, moving a rolling 7-amino acid wide window along the sequence of proteins with two or more highly stabilizing mutations, we computed the degron probability (using the random forest classifier described above) of each window, hence generating a degron probability profile of the entire protein sequence (panel 2 of Fig. 4a). We then identified peaks in the profile that may correspond to regions susceptible to allocate de *novo* degrons. (Note that because the approach is unable to identify the limits of a degron with certainty, some of the regions we identify may be longer than the average length of known degrons.) We retained as potential *de novo* degron any peak in the profile with at least two highly stabilizing mutations. We also required that the highly stabilizing mutations in the putative region caused on average a greater change of stability than other mutations in the protein (i.e., Z-score --computed using all the mutations observed in the protein as background-- higher than 0.5). Each *de novo* degron is thus characterized by two values: the mean stability change triggered by its mutations, and its probability, which we use to represent them in a two-dimensional graph (panel 4 of Fig. 4a). In this representation, *de novo* degrons are colored following the average Z-score of their high-impacting mutations (panel 3 of Fig. 3a).

**Figure 4.**
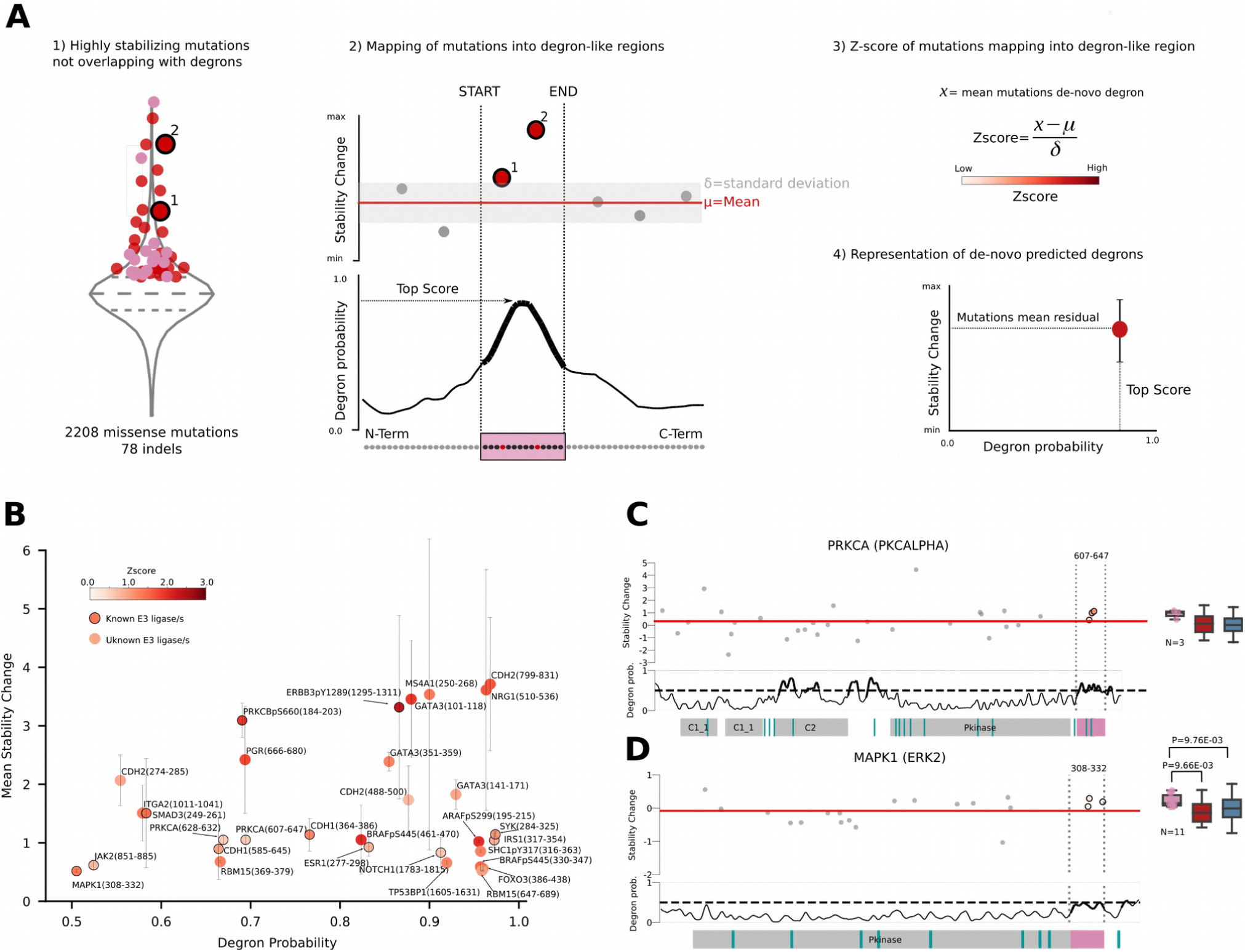
Identification of new instances of *de novo* degron instances. (a) Workflow to identify degron instances *de novo* from mutations in primary tumors. Highly stabilizing mutations that do not affect annotated or newly identified degrons are selected (panel 1). A profile of degron probability is computed across the full sequence of proteins bearing them (panel 2). Protein regions with degron probability above 0.5 affected by highly stabilizing mutations (positive Z-score among all mutations of the protein; panel 3) in at least two tumors are potential *de novo* degron instances (panel 4). (b) *De novo* degron instances (circles) represented in a plane of their classifier probability vs the mean stability change (with standard deviation as whiskers) of the mutations affecting them. Circle colors represent the Z-score of the stability change elicited by mutations. (c, d) Examples of *de novo* degrons identified in PRKCA (c) and MAPK1 (d). Top panels represent the mutations observed across tumors along the sequence of either protein. Dashed vertical lines delimit *de novo* degrons, with mutations within them colored following their elicited stability change. Bottom panels represent the profile of degron probability along the sequence of each protein. Colored bars below represent the location of protein domains (gray), annotated degrons (blue) and *de novo* degrons (pink). The boxplots correspond to the stability change in the three groups of proteins of the same color as Figures 3b-g.

To test the validity of this approach we applied it to the three annotated degrons with two or more highly-stabilizing mutations (the βTRCP inTRCP degron of CTNNB1 and the two Kelch-KEAP1 degrons of NFE2L2). We were able to clearly identify two of them as stretches of the protein sequence with a peak of degron probability above 0.5 (Figs. S4a,b). One of the Kelch-KEAP1 degrons is scored just below this threshold (at 0.48 degron probability). These results demonstrate that the approach described above is useful to specifically identify *de novo* degrons bearing recurrent highly-stabilizing mutations.

We found 31 regions of 23 proteins containing *de novo* degron sequences (Fig. 4b), for 13 of which there is evidence of direct interaction with at least one E3 (circles with border). Some of these *de novo* degrons were recurrently (in more than two samples) mutated across primary tumors. This is the case, for example, of the protein kinase C alpha (PRKCA; Fig. 4c) and beta (PRKCB; Fig. S4c), where we identified *de novo* degrons in regions spanning between amino acid residues 277 and 298, and between residues 184 and 203, respectively. The *de novo* degron identified in PRKCA is affected by 3 non-synonymous mutations, which trigger a stabilization of the protein with respect to its wild-type species. PRKCA is a known target of at least two E3s: a complex integrated by RBCK1 and RNF31 (Nakamura et al., 2006), and TRIM41 (Chen et al., 2007).

We identified another *de novo* degron in a region located between amino acid residues 308 and 332 of the protein product of MAPK1 (Fig. 4d). MAPK1 is known to be activated and ubiquitinated by MAP3K1, a protein with dual kinase and E3 ligase activity, in response to several stimuli. Aspartate residues at positions 316 and 319 of MAPK1 (within the *de novo* degron) are known to be important for its interaction with MAP3K1, both for phosphorylation and ubiquitination (Lu et al., 2002), but the degron remained unknown. Other *de novo* degrons (Figs S4d-g) include those of ERBB3, ESR1, ARAF and BRAF. Although the protein product of ERBB3 is known to be the target of the RNF41 E3 ligase (Carraway, 2010), its specific degron is unknown. The regulation of the half life of ESR1, encoding the estrogen receptor alpha is very complex, and includes the participation of the HSP90 and CHIP and other players (Berry et al., 2008; Duong et al., 2007). ARAF, one of the three Ras-binding proteins in the Ras-MAPK pathway is known to interact with the E3 HERC2 (Galligan et al., 2015). Finally, we identified two *de novo* degrons in BRAF flanking an annotated SCF_FBW7_1 degron, which recent studies suggest is not the sole responsible for the regulation of BRAF abundance (Hernandez et al., 2016).

### Degron-affecting mutations are positively selected in tumorigenesis

With the comprehensive proteome-wide catalog of potential new degrons (new instances of known degrons and *de novo* degrons), we proceeded to assess their involvement in carcinogenesis upon mutations across tumor types. Deviations of the observed mutations across tumors from their expected null pattern are routinely exploited as signals of positive selection by methods aimed at identifying cancer driver genes (Bailey et al., 2018; Gonzalez-Perez et al., 2013a; Tamborero et al., 2013). We inferred that the principles underlying these methods could be adapted to identify degrons under positive selection in tumorigenesis. Hence, we developed two methods to detect degrons whose mutational pattern deviates from the expected given the null mutational model.

The first method (SMDeg) probes the overrepresentation of missense mutations in new instances of known degrons or *de novo* degrons with respect to their number inferred from the distribution of all missense mutations observed in the protein. Briefly, SMDeg first counts the number of missense mutations observed in a degron (and 11 upstream and downstream flanking amino acids) and outside across tumors (six and two mutations, respectively in the toy example in the top panel of Fig. 5a). These eight non-syonymous mutations are then randomly re-sampled 1000 times along the nucleotide sequence of the gene, following the probability of mutation of each base, derived from its tri-nucleotide context (Mularoni et al., 2016) (Fig. 5a, middle panel). Finally, the observed and average number of mutations in the degron and outside of it are compared using a G-test of goodness-of-fit (bottom panel of Fig. 5a), from which the SMDeg p-value is derived. New instances of known degrons or *de novo* degrons with FDR < 10% (Benjamini-Hochberg) are deemed positively selected in tumorigenesis.

**Figure 5.**
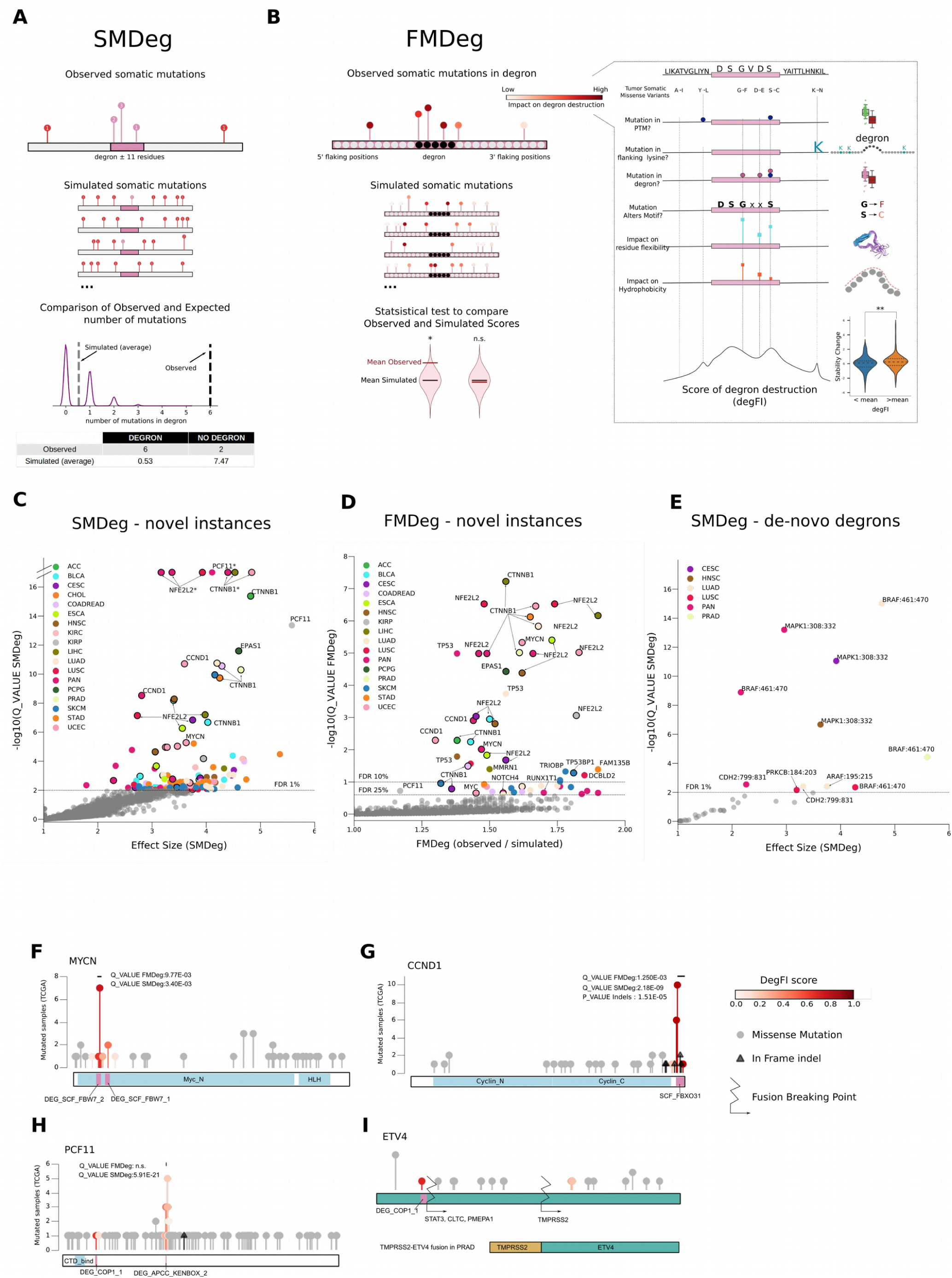
Degrons driving tumorigenesis. (a) Diagrammatic depiction of the SMDeg test. Top panel: observed mutations in a gene where a novel degron instance has been identified (pink rectangle). Middle panel: the same number of mutations are randomly resampled following the mutational probability of each nucleotide in the gene several times to produce their expected distribution. Bottom panel: the observed and expected number of mutations within and outside the degron are compared via a Gtest. (b) Diagrammatic depiction of the FMDeg test. Top panel: observed mutations in a novel degron instance (black positions) and its flanks (white positions) are scored using the degFI (a score to measure their impact on degron functionality; right panel). Middle panel: the same number of mutations are randomly sampled from all positions in the degron and its flanks following the mutational probability of each nucleotide to produce the expected distribution of degFI values. Bottom panel: the mean degFI of observed mutations is compared to the expected distribution of degFI values to yield an empirical p-value. (c, d) Right-tail volcano plot representing novel degron instances that appear significant (FDR < 1%) in the SMDeg (c), or significant (FDR < 10%) or nearly significant (FDR < 25%) in the FMDeg test (d) across cohorts of primary tumors (color-coded). The PAN cohort contains a union of all TCGA tumors. Annotated degrons appear circled in black. (e) *De novo* degrons that appear significant in the SMDeg (c) test across cohorts of primary tumors (color-coded). The PAN cohort contains a union of all TCGA tumors. (f-i) Needle-plots representing the distribution of mutations, in-frame indels and/or fusion breakpoints in the PAN cohort along the sequences of MYCN (f; significant in SMDeg and FMDeg), CCND1 (g; significant in SMDeg, FMDeg and the in-frame indels test), PCF11 (h; significant in SMDeg), and ETV4 (i; non-tested). The four genes harbor novel degron instances (purple rectangles).

The second test (FMDeg) computes the deviation of the average functional impact of missense mutations hitting the sequence of new instances of known degrons from their expected impact given the sequence composition of the degron match (left panel of Fig. 5b). To measure the functional impact of individual missense mutations --i.e., their potential impact on the capability of the degron instance to bind its cognate E3 and become ubiquitinated-- we designed an *ad hoc* score (degFI; right panel of Fig. 5b). The degFI is based on the structural features deemed most important to define a degron by the classifier described above. For example, it takes into account whether the mutation is projected to affect the sequence motif of the degron, its flanking phosphorylation sites, or its flexibility (see Methods). To assess the usefulness of degFI to measure the effect of mutations on degron functionality, we tested if mutations with high degFI tend to change protein stability. We show that the stability of proteins bearing mutant degrons with high degFI scores (i.e., above the mean of the global distribution) is significantly higher than that of proteins bearing mutations with degFI scores below the mean (Mann-Whitney one-side test p-value<0.01; boxplot of Fig. 5b right panel). (Note that since some of the features within the degFI, such as the motif conservation, can only be computed for new degron instances, the FMDeg cannot be applied to *de novo* degrons). FMDeg hence computes the average degFI of the mutations observed in each degron match, and the distribution of the average degFI from 10,000 samples of the same number of mutations in the matching sequence, drawn following the mutation probability of each nucleotide in the gene as described above for SMDeg. Finally, it derives an empirical p-value from the fraction of random samples with average degFI higher than (or equal to) the average degFI of observed mutations (Mularoni et al., 2016). New instances of known degrons with FDR < 10% (Benjamini-Hochberg) are deemed positively selected in tumorigenesis.

We applied both tests to the 18239 novel degron instances identified before. We also applied the SMDeg to the 31 *de novo* degrons identified in the previous section. We applied both methods in a pan-cancer manner and within each tumor cohort. (Note that in this case, we were able to use the 9550 TCGA non-hypermutated primary tumors with somatic mutations data; see Methods). The distribution of p-values produced by each was very similar to the null distribution, except for the few significant degrons, which showed the correct calibration of both methods (Fig. S5a,b). Five out of the 26 genes bearing degrons that are significant or nearly significant according to both tests (intersection) in the TCGA analysis were oncogenes. This is a significant overrepresentation with respect to the proportion of oncogenes among all tested genes (Fisher’s OR=5; p-value=0.01).

Figures 5c, d and e represent the array of significant degrons identified by each test, with annotated degrons circled in black (in Figs c and d). NFE2L2 and CTNNB1 annotated degrons are significant both across all tumors and across several tumor-type cohorts according to both methods. The annotated SCF_FBW7_2 degron in the known oncogene MYCN appears significant in both tests across the pan-cancer cohort (Fig. 5f). Interestingly a new instance of the APCC_KENBOX_2 degron in PCF11, a gene involved in the processing and maturation of mRNA, is also among the top ranking instances of both methods. Mutations in this novel degron instance, accumulate significantly more than expected (Fig. 5g). The novel FBXO31 degron instance of CCND1, which carries an abnormally high number of impacting missense mutations and in-frame indels (Fig. 5h), is identified as significant by both methods in uterine carcinomas and in the pan-cancer cohort. Both tests also identify the novel VHL degron instance of EPAS1 in pheochromocytomas and paragangliomas as highly significant.

A region likely to contain a *de novo* degron between amino acid residues 461 and 470 of BRAF (Figs. 4b and S4g) is identified as highly significant by the SMDeg test in several tumor types (Fig. 5e). It also identifies the MAPK1 region between residues 308 and 332, which likely contains a *de novo* degron of MAP3K1 (Figs. 4b and d) as a potential cancer driver (Fig. 5e).

An abnormal stabilization of oncoproteins may also be caused by in-frame indels (as in the cases of NFE2L2, CTNNB1 and CCND1 exemplified in Figures 3b, c and d, represented by pink triangles) or fusions that disrupt the sequence of their degrons. To identify degrons disrupted by in-frame indels more than expected by chance --as another signal of positive selection in tumorigenesis-- we carried out a simple overrepresentation probe (using Fisher’s exact test) for all degrons with probability higher than 0.5 and with at least 3 in-frame indels. The degrons of NFE2L2, CTNNB1 and CCND1 (and others) are significantly enriched for in-frame indels (Table S4), thus identifying a potential novel mechanism of oncogenesis that involves their abnormal stabilization via disruption of their ubiquitin-mediated degradation. We reasoned that degrons may also be disrupted by fusion events. For example, a COP1_1 degron close to the N-terminal of ETV4 is abrogated via fusions of the gene with several partners in 4 tumors (Fig. 5i). Two different breakpoints produce a fusion product that lacks this degron, although their functional effects remain to be determined. We thus compiled a list of 130 proteins whose degrons are disrupted by fusion events in at least two primary tumors in the cohort (Table S4).

Several degrons deemed significant in the SMDeg and FMDeg tests across primary tumors also exhibit signals of positive selection across the 899 cancer cell lines in the CCLE (Fig. S5c,d). The degrons in CTNNB1, NFE2L2 and CCND1 are amongst these. Some novel interesting candidates, such as ETV5 (a paralog of ETV4 with a novel instance of the SCF_FBW7_1 degron significant in SMDeg; Fig. S5e), CCND3 (a paralog of CCND1 with an instance of FBXO31 degron significant in both SMDeg and FMDeg; Fig. S5f), and USP36 (a protein involved in RNA maturation with a recurrently mutated instance of SPOP_SBC_1 degron; Fig. S5g) are identified as well.

In summary, we uncovered 49 cancer driver degrons (either significant in both the SMDeg test and FMDeg tests or significant in the in-frame indels tests) across 32 cancer types (Table S4) and 899 cancer cell lines. Our results show that the contribution of degron destruction to tumorigenesis goes far beyond the few known examples of mutations that destroy degrons of oncoproteins (Guharoy et al., 2016; Mészáros et al., 2017).

### The downstream effect of mutations of driver E3s

Alterations affecting E3s also contribute to the role of UPS dysregulation in tumorigenesis. Outside the well-known cases of E3-target links in which the degron is known, the immediate downstream effects of E3 alterations (i.e., whose target oncoproteins become abnormally stabilized) are unknown. We reasoned that newly identified degrons and the measurement of protein stability across tumors provided a way to thoroughly explore this question for the first time.

We started by detecting E3s with signals of positive selection in tumorigenesis. To that end, we employed dNdScv (Martincorena et al., 2017) and OncodriveFML (Mularoni et al., 2016). Any E3 exhibiting significant recurrence of non-synonymous mutations (detected by dNdScv) or significant bias towards high impacting mutations (FMbias; probed by OncodriveFML) either in a cohort of a particular cancer type or across the pan-cancer cohort was considered a cancer driver E3 (Fig. 6a and Table S5). Several E3s, such as FBXW7, NFE2L2, SPOP, APC, KEAP1, MAP3K1, VHL, and RNF43 appear significantly mutated or FMbiased in more than one cancer type. Our results are similar to a list of recently published driver E3s across TCGA primary tumors (Ge et al., 2018). We also identified driver E3s across cancer cell lines (Fig. S6 and Table S5).

**Figure 6.**
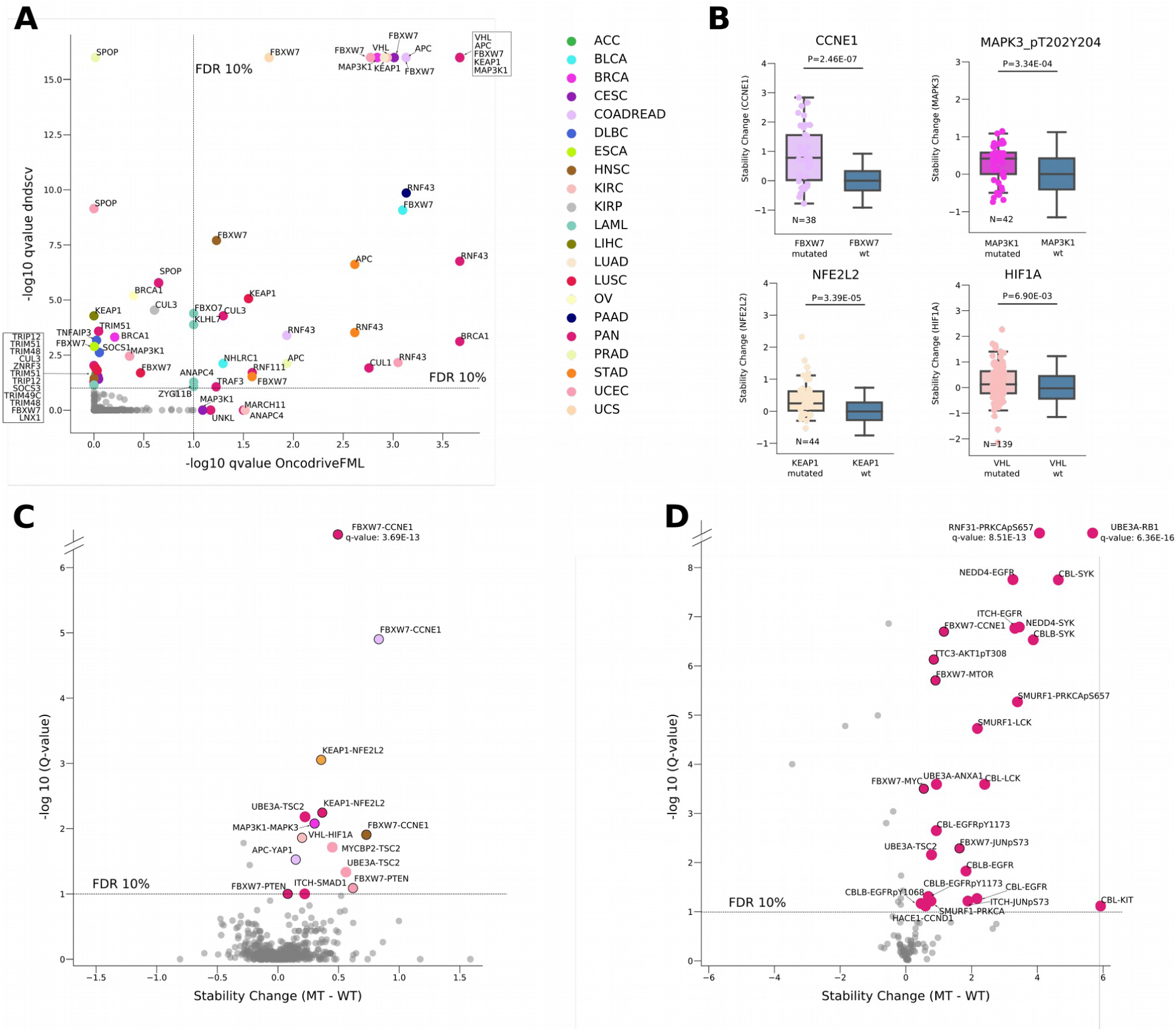
Downstream effects of driver E3 alterations. (a) Driver E3s across cohorts of different cancer types (color-coded) are identified through signals of positive selection detected by OncodriveFML (Mularoni et al., 2016) and dNdScv (Martincorena et al., 2017). (b) Clockwise from top-left corner, the stability of CCNE1, MAPK3 (detected through an antibody against a di-phosphorylated form), NFE2L2 and HIF1A significantly increase upon mutations of FBXW7, MP3K1, KEAP1 and VHL, respectively. (c) Changes of protein stability elicited by mutations of all driver E3s across cohorts of different primary tumor types. (d) Changes of protein stability elicited by mutations of all driver E3s across cancer cell lines.

We then explored the downstream effects of mutations in driver E3s on the stability of proteins bearing instances of their cognate degrons, either annotated or newly identified. To this end, we employed the approach introduced before to evaluate the effect of mutations on protein stability, i.e., compare the stability of all proteins in tumors harboring a mutant E3 and tumors with its wild-type form. (Tumors harboring mutations or high-level CNAs of the E3 target were filtered out from this analysis; see Methods.) For example, we found that the stability of CCNE1 is significantly higher in colorectal tumors bearing FBXW7 mutations than their counterparts with the wild-type E3 (Fig. 6b, top left panel). Although the FBXW7 degron of CCNE1 is only rarely mutated in tumors (4 mutated samples in TCGA), the gene itself is amplified, producing an overexpression of the protein akin to that caused by the loss-of-function mutation of its E3. Analogously, the stability of MAPK3, NFE2L2 and HIF1A significantly increases in breast tumors, lung adenocarcinomas and kidney clear cell carcinomas carrying mutations of their cognate E3s, respectively (Fig. 6b, other panels).

The systematic application of this test to all proteins with RPPA information probing all driver E3s across every TCGA cohort revealed several significantly stabilized targets (Fig. 6c). E3-target pairs that are known to interact, and which therefore constitute cases of more likely direct stabilization, are marked with an outer circle in the figure. This stabilization analysis picks up some well-known E3-target relationships (e.g., FBXW7-CCNE1 and KEAP1-NFE2L2), but also highlights novel interesting links. For example, the potential ubiquitination of MAPK1 by MAP3K1 through a *de novo* degron, which we discussed above, is now given more credence by the discovery that mutations of MAP3K1 in breast tumors significantly increase the stability of MAPK3, a paralog of MAPK1 with high sequence identity. TSC2 is significantly more stable in breast and uterine carcinomas with mutations of UBE3A. Other interesting cases are uncovered by a similar analysis across cancer cell lines (Fig. 6c). PRKCA, a protein in which we identified a *de novo* degron is significantly stabilized upon mutations of two E3s, RNF31 and SMURF1. In both cases, the increase in stability is more notorious for a phosphorylated form of the protein (in serine 657), that is 10 amino acids to the C-terminal of the *de novo* degron (Fig. 5c).

Summing up, alterations affecting driver E3s (Table S5) significantly increase the stability of the proteins carrying their cognate degrons.

### The disruption of UPS is an important contributor to tumorigenesis

The tumorigenic function of oncoproteins may be achieved through both the increase of their activity or the number of their units in the cell (Fig. 7a). Different types of alterations are known to produce the latter. For example, the increase in the number of genomic copies of an oncogene in a tumor (such as MYC in breast adenocarcinomas), or its fusion to another gene (e.g., BRAF in piloastrocytomas) result in the overexpression of its protein product. Alterations that disrupt its targeted degradation --either affecting its cognate E3 or its degron-- produce similar outcome (Fig. 7b). It is possible to exemplify the variety of alterations leading to increased number of protein units through the case of CCNE1 (Fig. 7c). The level of its protein product is higher in tumors bearing either a mutation in its degron, an amplification of the gene, and/or a mutation affecting FBXW7 (Fig. 7c) than in tumors with neither of these alterations. Interestingly, in tumors with concurrent amplification of CCNE1 and mutation of FBXW7 the level of the protein product is higher than in those bearing either alteration. Overall, almost 10% of all tumors in the TCGA pan-cancer cohort carry at least one alteration that results in the increase of CCNE1 protein levels.

**Figure 7.**
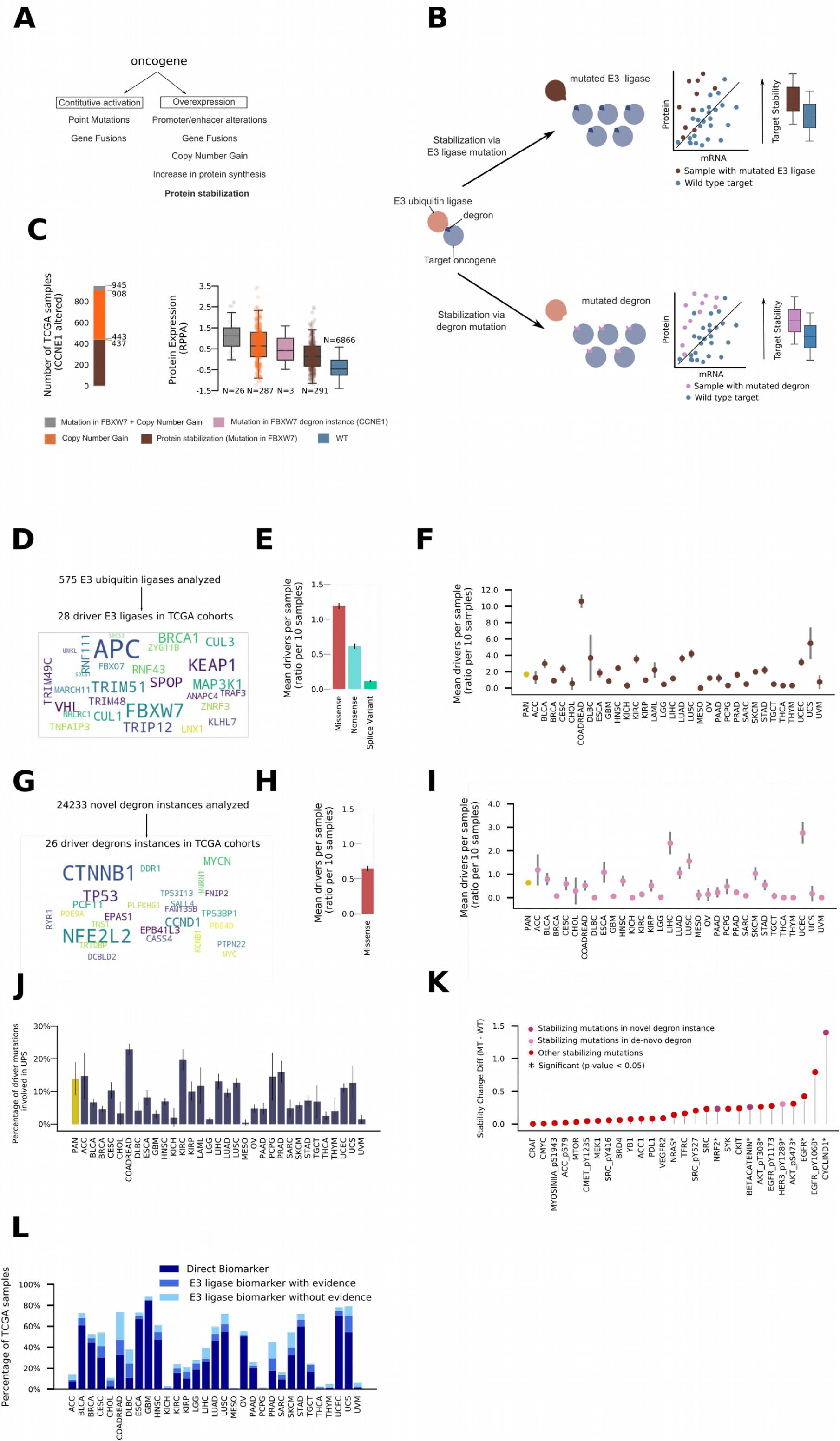
Contribution of UPS to the emergence of primary tumors. (a) Types of tumorigenic alterations of oncoproteins. (b) Types of alterations of the UPS that may result in abnormally high levels of oncoproteins. (c) Different alterations that elicit abnormally high levels of the CCNE1 protein product. Left bar: number (and proportion) of TCGA primary tumors affected by either alteration type. Right boxplot: levels of CCNE1 protein product in tumors bearing different alteration types. (d) Mutation frequency of driver E3s represented as a wordcloud. (e) Average number of driver mutations (with different consequence type) in driver E3s per 10 TCGA tumors. (f) Average number of driver mutations (sum over consequence types) in driver E3s per 10 tumors in different TCGA cohorts. (g) Mutation frequency of driver novel degron instances represented as a wordcloud (of their proteins). (h) Average number of driver non-synonymous mutations in driver novel degron instances per 10 TCGA tumors. (i) Average number of driver non-synonymous mutations in driver novel degron instances per 10 tumors in different TCGA cohorts. (j) Proportion of driver mutations (in 405 known cancer genes) that are contributed by UPS in different TCGA cohorts. (k) Top 28 most stabilized proteins upon mutations in novel degron instances (dark purple), *de novo* degrons (light purple), or in other sites in the sequence (red). (l) Opportunities for repurposing anti-cancer drugs used in the clinic to target the abnormal increase of oncoproteins levels to tumors bearing loss-of-function alterations of their cognate E3s.

Next, we asked how much alterations of the UPS across cancer types contribute to tumorigenesis. To tackle this problem, we used dNdScv (Martincorena et al., 2017), through which we are able to estimate the number of non synonymous mutations in any set of genes across a cohort of tumors that exceeds the expected under neutrality (after correcting for the mutational processes and the regional covariates of neutral mutation rate). We assume that these non synonymous mutations in excess represent the positively selected drivers of the tumors in the cohort (Martincorena et al., 2017). We can subsequently estimate the average number of driver mutations in each tumor of the cohort (see Methods). We thus broke down the aforementioned question to compute the contribution to tumorigenesis of driver E3s and degrons. For the former, we focused on the 28 E3s with at least one signal of positive selection across any cohort of primary tumors in TCGA (Fig. 7d). We computed the excess of non-synonymous, nonsense, and splice-affecting mutations in the set of these driver E3s across the pan-cancer cohort (Fig. 7e), and their combined contribution to the excess mutations in each cohort (Fig. 7f). On average, almost 2 in 10 primary tumors in TCGA carry a driver mutation affecting an E3; the majority of these driver mutations are non-synonymous. The contribution in some cohorts is greater, with all colorectal tumors carrying --on average-- more than one driver mutation of an E3, the vast majority of them, mutations of APC and RNF43. The collective contribution of the 26 driver degrons (two signals of positive selection across any TCGA cohort; FDR < 25%) to tumorigenesis is smaller (Fig. 7g). On average, almost one in 15 primary tumors carries a non-synonymous driver mutation affecting one of these degrons (Figs. 7h, i). There are again instances of greater contribution across specific cohorts, such as the βTRCP inTRCP degron of CTNNB1, with a driver mutation in one of four liver carcinomas.

To further understand the relevance of this contribution of UPS mutations (UPS-drivers) to tumorigenesis, we compared it with the driver mutations contributed by a list of 369 well-known driver genes (Futreal et al., 2004), to which we added driver E3s and driver degrons. Overall, these cancer genes contribute close to two driver (all-drivers) mutations per tumor, which makes the ratio of UPS-drivers-to-all-drivers is close to 0.13. In other words, 1 in 7 driver mutations attributable to well known cancer genes is contributed by UPS mutations. Still, due to the scarcity in the identification of degrons, these results are very likely an underestimation of the contribution of UPS to tumorigenesis via mutations. One clue that this is probably the case is the fact that some of the most highly stabilizing mutations (amongst the 209 with protein levels information) do not occur in known or newly identified degrons (Fig. 7k).

The identification of the oncogenic downstream targets of E3 ligases expands the scope of genomic-driven anti-cancer therapeutic approaches. In effect, tumors driven by loss-of-function alterations of E3 ligases could in principle be targeted via inhibitors of their abnormally overabundant downstream targets. We computed the number of patients in TCGA cohorts that could benefit in principle of this “indirect” repurposing of anti-cancer drugs in clinical or pre-clinical stages (Fig. 7l). In each TCGA cohort we count the number of tumors that bear mutations affecting known E3 ligase targets that constitute known biomarkers of anti-cancer drug response obtained from the Cancer Genome Interpreter (Tamborero et al., 2018) (labeled as Direct Biomarker). For example, 67 (13%) TCGA uterine adenocarcinoma samples carry CCNE1 amplification, which constitute a pre-clinical stage biomarker of response to CDK2 inhibitors (Fig. S7). We also compute the number of tumors with alterations of the corresponding E3 ligases, the mutations of which increase the stability of the target protein (labeled as E3 ligase biomarker with evidence). Following upon the previous example, we find 27 (5%) TCGA uterine adenocarcinoma samples with FBXW7 mutations resulting in the increase of CCNE1 stability, which could in principle be targeted using the same CDK2 inhibitors. Finally, if we add all tumors with mutations of the E3 ligase without evidence of increased stability of the target (labeled as E3 ligase biomarker without evidence) other 42 (8%) FBXW7-mutant uterine adenocarcinoma samples could potentially be targeted using these drugs. These examples of repurposing anti-cancer drugs to target loss-of-function mutations of E3s could potentially benefit important fractions of tumors in certain cohorts, such as colorectal adenocarcinomas, which include 77 (14%) samples with E3 ligase biomarker with evidence and 143 (26%) samples with E3 ligase biomarker without evidence (Table S6).

## Discussion

Identifying the elements that integrate the UPS is key to understanding the operation of the turnover of proteins in the cell, the regulation of key processes, such as the cell cycle, and the role that their dysregulation plays in disease (Bassermann et al., 2014; Mészáros et al., 2017; Vu and Sakamoto, 2000). Here, we systematically identified new instances of known degrons across human proteins, and we provide evidences of the functionality of some of them. Such evidences encompass i) the abnormal stabilization of proteins upon mutations affecting their potential degrons; ii) signals of positive selection of the destruction of potential degrons in their pattern of mutations across tumors; and iii) the disruption of the function of these potential degrons, again resulting in the anormal stabilization of proteins upon mutations of the E3 ligases that “read” them. Along the manuscript, we highlighted some exemplary matches of degrons supported by these evidences; the complete list of matches, thoroughly annotated, and provided as Supplemental Table 2 is one of the main outcomes of our work (see below).

We acknowledge the limitations of the approaches taken in this study that emanate primarily from constraints of the data. On the one hand, only few instances of few degrons are known, which limits the space of exploration of new degrons; on the other hand, simultaneous measurement of genetic variants and the abundance of proteins and mRNAs in the same pool of cells is still rare. Moreover, while the whole exome and transcriptome can be readily probed across thousands of samples, the detection of protein abundance is still limited to either a small set of proteins across many samples (mostly using RPPA), or to many species across a limited number of samples (via MS). While the genetic variants analyzed may produce the observed effect through other mechanisms, such as increase of protein abundance via disruption of miRNA binding sites (Hausser et al., 2013) or change in translation efficiency, this problem is minimized by analyzing only coding mutations in potential degrons. Another potential caveat arises from the fact that in cancer cells --our experimental setting-- other parts of the UPS may be altered besides the specific within-degron variant under analysis, thus complicating the attribution of causality of the effect on protein abundance. Again we minimize this problem by removing samples carrying potentially obscuring alterations of known parts of the UPS (E2-enzymes, E3 ligases, adaptors, and deubiquitinases) and the protein under observation (amplifications or deletions). We envision that as more data of protein abundance --foreseeably through application of MS to both normal and cancer cells-- become available (as expected, for example from the enhanced GTEx consortium) (eGTEx Project et al., 2017), the application of the approach presented here will result in the discovery of yet unknown degrons in a true proteome-wide manner.

This work touches on key aspects of basic and cancer biology, and consequently, its important outcomes are of interest to several research communities. The list of matches of known degrons, the rationale and annotations used to prioritize them, and the shortlist of high-confidence matches constitute a valuable resource to UPS researchers and protein engineers. We have introduced a simple and novel framework for the analysis of the influence of any type of alterations (in *cis* or *trans*) on the stability of a given protein, and we have demonstrated its use with RPPA and MS protein abundance data. We also presented a score of functional impact of mutations in degrons based on the disruption of their defining sequential and structural properties. Finally, we introduce the first comprehensive landscape of UPS disruption in tumorigenesis, including assessing the effect of alterations in E3 ligases and their targets. Our results shed light on the cellular downstream effects of a specific subset of driver mutations (affecting UPS), a key line of research to bridge the gap between cancer genomics and cancer personalized medicine (Gonzalez-Perez and A, 2016; Gonzalez-Perez et al., 2013b, 2016). We demonstrate the usefulness of such research line exploring the landscape of the opportunities to repurpose anti-cancer drugs to target loss-of-function alterations of E3s.

Our results provide the first systematic assessment of the contribution of alterations of the UPS machinery to tumorigenesis. We estimated that 1 in 7 driver mutations attributable to well known cancer genes is contributed by elements of the UPS. Therefore, alterations of the UPS that result in the abnormal stabilization of oncoproteins constitute a frequent mechanism of tumorigenesis. While we also provided clues that this assessment represents an underestimation, we foresee that the approaches developed and tested here will pave the way to uncover a more complete landscape of the involvement of UPS in tumorigenesis in the near future.

## METHODS

### Data collection and preprocessing

#### TCGA data

Level 4 cohort-specific RPPA data was downloaded from the TCPA portal (http://tcpaportal.org/tcpa/download.html; version 4.2; 07/02/2017). The dataset included 7,663 samples and 258 different antibodies recognizing to 209 proteins. Epitope information was manually curated from the TCPA portal and the vendor’s website. Antibodies without vendor’s information were excluded from the study.

Cohort-specific RNASeq data was downloaded from the TCGA Firebrowse portal (http://firebrowse.org/; 21/07/2016). The RNASeq dataset included 10,0082 samples.

Cohort-specific CNA data was downloaded from the TCGA Firebrowse portal (http://firebrowse.org/; 14/12/2016). The CNA dataset included 10,842 samples.

Somatic mutations were downloaded from the TCGA Firebrowse portal (http://firebrowse.org/; 29/12/2016). Somatic mutations were filtered using the column FILTER from the TCGA calling pipeline (i.e., only mutations with column FILTER == ‘PASS’ were kept). Multiple alterations within the same gene in one particular sample were aggregated. Each sample’s phenotype was defined as the first hit from the following list of consequence types: Splice_Site, Nonsense_Mutation, Frame_Shift_Del, Frame_Shift_Ins, Nonstop_Mutation, Translation_Start_Syte, In_Frame_Del, In_Frame_Ins, Missense_Mutation, Intron, Silent, 5’UTR, 3’UTR, IGR, 5’Flank, 3’Flank. The somatic mutations dataset included 9,007 samples.

Colon adenocarcinoma (COAD) and rectum adenocarcinoma (READ) cancer types were merged under the label COADREAD in all sources of data.

#### Cancer Cell Line Encyclopedia data (CCLE)

Cell line specific RPPA (CCLE_RPPA_20180123; 10/10/2018), Antibody information (CCLE_RPPA_Ab_info_20180123; 10/10/2018), RNA (CCLE_DepMap_18q3_RNAseq_RPKM_20180718; 10/10/2018), CNA (CCLE_copynumber_byGene_2013-12-03; 10/10/2018) and somatic mutations (CCLE_DepMap_18q3_maf_20180718; 10/10/2018) were downloaded from the Broad Portal (https://portals.broadinstitute.org/ccle/data). Antibodies not matching with the RPPA identifier were manually updated. The final dataset included 214 antibodies in 899 cell lines. RPKM were transformed into log2(RPKM). CNA raw data was discretized into 5 levels (i.e, high-level deletion, low-level deletion, diploid, low-level amplification and high-level amplification) using cell line specific bounds form GISTIC2 (CCLE_copynumber_2012-04-05.seg.sample_cutoffs.txt). For each cell line if the raw CNA was greater than the upper bound, then the CNA it was considered a high-level amplification. Similarly, raw CNAs below lower bounds were considered high-level deletions. Cell lines with raw CNA greater than 0.3 and raw CNA lower than upper threshold were considered low-level amplifications. Cell lines with raw CNA greater than −0.3 and raw CNA lower than lower bound were considered low-level deletions. Cell lines with raw CNA *≤* 0.3 and CNA *≥* −0.3 were therefore considered as diploid. Finally, when there was no information available about lower or upper bounds, the average value across all cell lines was used used as cutoff. Somatic mutations were processed following the TCGA protocol and protein position was parsed to include information about the aminoacid harbouring the somatic alteration.

#### Mass Spectrometry data

Mass-spectrometry (MS) data from human high-grade serous ovarian cancer (OV) (Zhang et al., 2016) and breast cancer (BRCA) (Mertins et al., 2016) TCGA cohorts was downloaded from the Clinical Proteomics Tumor Analysis Consortium using TCGA-assembler 2 (Wei et al., 2018). The MS dataset contained 280 tumor samples, including 105 BRCA and 175 OV samples, and a total of 11,064 proteins with their level measured through mass-spectrometry.

#### Curated list of proteins involved in ubiquitination

The list of proteins involved in ubiquitination (UBSs) and deubiquitination (DUBs) was manually created by integrating prior knowledge from UniProt (Bateman et al., 2017) and E3NET (Han et al., 2012). The final list of proteins involved in ubiquitination included 977 identifiers of human proteins.

#### Curated list of E3 ligase substrate interactions

A curated list of E3 ligases and their degradation substrates was downloaded from (http://pnet.kaist.ac.kr/e3net/; 17/10/2018) (Han et al., 2012). Non-human interactions were filtered out. The APC-CTNNB1 interaction was manually added to the list. The final list of proteins involved in ubiquitination included 833 pairs of interactions between E3 ligases and human substrates.

#### List of protein-protein interactions from STRING

We downloaded human protein-protein interaction data from STRING (Franceschini et al., 2013) (<u>9606.protein.links.detailed.v10.5.txt.gz</u>; 19/02/2018). STRING identifiers were mapped to HUGO symbols using UniProt. Interactions that do not include a protein involved in ubiquitination (see Curated list of proteins involved in ubiquitination) were discarded. Similarly, pairs with an STRING score below 300 were filtered out. The final list of 2,568,513 protein-protein interactions involved 18,403 proteins, including 566 E3 ligases.

#### Combination of data for TCGA and CCLE

We merged the information (i.e., RPPA or MS Log_Ratio(iTRAQ), RNA, CNA and somatic mutations) available for each gene (represented by a HUGO symbol) in each TCGA primary tumor or CCLE cell line. We then annotated, for each pair, whether an upstream E3 ligase of the protein harbours a non-synonymous variant in that sample/cell line. We used the curated dataset from (Curated list of E3 ligase substrate interactions) as source of curated interactions. Additionally, for each sample-protein pair with mutations in the sample we annotated whether the mutation falls into the epitope region of the antibody. Finally, samples/cell lines with non finite values of RPPA were discarded.

The matched RPPA TCGA dataset contained 6,907 samples and 236 antibodies that recognize 193 proteins.

The matched RPPA CCLE dataset contained 861 samples and 200 antibodies that recognize 157 proteins.

The matched MS TCGA dataset contained 210 samples and 10,065 proteins.

## Protein-mRNA regression

### Calculation of raw residuals

#### TCGA

Given an antibody-tumor type pair in the TCGA dataset, the relationship between the protein level --measured by the RPPA-- and the mRNA --measured by the log2(RPKM)-- was estimated by means of a robust linear regression approach, as we wanted to derive a linear model insensitive to outliers and high leverage points, the fitting method of choice being Iteratively Reweighted Least Squares (IRLS). Samples with mutations, high level amplifications (i.e., CNA *≥* 2), or alterations in annotated upstream E3 ligases were not included in the calculation of the regression. For each gene in each sample, the residual, defined as the y axis distance from the protein RPPA value to the regression line given its mRNA, was calculated and then normalized (see residual normalization and Supplemental Note 1 for further information about the normalization). Mass spectrometry cohorts followed similar protocol where the Log_Ratio(iTRAQ) measured protein expression.

#### CCLE

Due to the inability to group samples by cohorts in the CCLE dataset, all cell lines were joined to perform the robust linear regression by IRLS. However, this grouping may result into heterogeneous distribution of mRNA-protein abundance due to tissue-specific expression patterns. More specifically, we observed that some cell lines displayed a bimodal distribution consistent with low versus high expression profiles. Figure Supplemental Note 1 illustrates this particularity thought the example of CDH1. Figure Supplemental Note 1a shows that CDH1 has a bimodal distribution across CCLE cell lines. To address this issue, each antibody was first classified into unimodally or bimodally distributed. This was done by fitting a two-dimensional Gaussian mixture model of the joint distribution of mRNA and RPPA. Antibodies whose distribution were more accurately fit by two components were classified as bimodal. The thresholds defined to classify as bimodal were Bayesian Information Criterion (BIC) difference between the models (i.e., the differences in the BIC score of the model with one component minus the bimodal model) greater than 150 and euclidean distance between centroids of the model with two distributions greater than For antibodies classified as bimodally distributed, the Gaussian mixture model assigned to each cell line the closest distribution. Figure Supplemental Note 1b represents the classification of cell lines into two components according to the their RPPA and mRNA value for the CDH1 antibody. Finally, for both unimodals and the two independent distributions of bimodal antibodies, robust regression was performed as previously described. Figure Supplemental Note 1c represents the robust regression on the two independent distributions of CDH1 across CCLE cell lines. Samples with somatic mutations, high-level amplifications or alterations in upstream E3 ligases were not considered in the classification nor the example of CDH1. Scikit-learn package of python was used to perform the classification.

### Residuals normalization

In order to render the residuals comparable across protein-tumor types it is necessary to correct for the influence that the dispersion of the samples’ protein and mRNA levels have in the slope of the robust regression line which the residuals are derived from. Hereto we describe the normalization procedure, with examples that illustrate its necessity.

Residual values were first normalized dividing each observed raw residual value by the cohort’s RPPA -or Log_Ratio(iTRAQ)-standard deviation (std). Example 1 (Supplemental Note 1a,b,c) justifies the need to apply such normalization. In this example, two pairs of protein-cancer types (EPPK1-COADREAD and CDK1-PRAD) with similar log2(RPKM) dispersion showed distinct RPPA dispersion (EPPK1 std=1.13; CDK1 std =0.04). Such differences have an influence on the distribution of raw residuals (Supplemental Note 1c, left panel). After dividing the raw residual by the RPPA std, both cases show similar distribution of residuals (Supplemental Note 1c mid and right panels).

Likewise, Example 2 (Supplemental Note 1d,e) justifies the need to normalize by mRNA expression. Lower mRNA dispersions will always tend to make higher regression slopes, thereby pushing the residuals of the mRNA outliers up. In Example 2, the first pair, PGR-BRCA (Supplemental Note 1d), shows a linear correlation between mRNA expression and RPPA (R=0.76; p-value∼0), which likely represents a biological effect. The second pair, ANXA7 in PRAD (Supplemental Note 1e), also shows linear correlation (R=0.34, p-value=1.8e-10), although it is driven by outliers and does not necessary represent mRNA-RPPA linear correlation. To rule out distortion of the regression slope by unbalanced dispersions between the axes, we rescaled both quantities by their respective standard deviations:

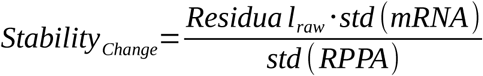

## Degrons

### Human proteome sequences

Amino acid sequences from reviewed 32,022 human isoforms were downloaded from UniProt (Bateman et al., 2017). Each sequence was associated with a UniProt isoform id and a HUGO symbol.

### List of annotated degrons

Degron motifs (i.e., consensus motifs of a particular E3 ligase/adaptor) and degron instances in the human proteome were downloaded from ELM (Dinkel et al., 2016) (http://elm.eu.org/downloads.html; 21/03/2017). Motifs not directly involved in ubiquitination were filtered out. In this study we focused on internal motifs, therefore motifs recognizing N or C-terminals were not considered. The lists were updated with information recorded in (Mészáros et al., 2017). Degron motif classes was manually curated from Table 1 and instances from Supplemental Table 1. Motif instances not matching with the UniProt canonical sequence were filtered out. The final tables (Supplemental Table 1) of degron motifs and instances included 29 ubiquitination motifs and 150 instances in 98 different proteins.

### Calculation of biochemical features

Here we specify the software and parameters that were used to compute the biochemical features of degrons:

- Disorder (DISORDER). IUPRED software (Dosztányi et al., 2005)was used to calculate protein disorder. Short disorder was chosen as prediction type. Degron instance disorder was calculated as the mean value of the motif amino acids.
- Solvent accessibility and secondary structure (S.A., COIL, HELIX and SHEET). SPIDER2 tool (Yang et al., 2017) with default parameters was used to calculate residue specific solvent accessibility alongside its secondary structure. Degron solvent accessibility and secondary structure specific probability (i.e., coil probability, α-helix probability and βTRCP in-sheet probability) were calculated as the mean value of the motif residues.
- Rigidity (RIGIDITY). Dynamine (Cilia et al., 2013) software with default parameters was used to calculate residue-specific flexibility. Degron flexibility was calculated as the mean value of the motif aminoacids.
- Stabilization upon binding (ANCHOR). ANCHOR program (Mészáros et al., 2009) with default parameters was used to calculate the anchoring score of each degron. Degron anchoring potential was calculated as the mean value of the motif aminoacids.
- Flanking conservation (FCONS). First, for each input sequence multiple sequence alignment of orthologs proteins were calculated using the Gopher tool from Bioware (http://bioware.ucd.ie/∼compass/biowareweb/Server_pages/gopher.php) [PMID: <u>16556315</u>]. Next, residue-specific conservation score was calculated using the Capra JA. and Singh M. method *(Capra and Singh, 2007)* scoring=shannong_entropy; gap_cutoff=0.9; matrix=blosum62). Finally, for each degron, the average conservation of degron residues was divided by the average conservation of flanking amino acids (eleven residues upstream and eleven amino acids downstream).
- Structured domains (DOMAIN). Information of protein domains was downloaded from Pfam (Finn et al., 2016)(09/05/2017). Degrons with at least one residue overlapping with a Pfam domain were labelled as degron domains. Therefore, domain-free domains did not overlap with any Pfam domain.
- Number of degron-associated phosphorylation sites (PHOSPHO_SITES). The degrons flanking sequences were scanned searching for known phosphorylation sites (see below). The count represented the number of phosphorylated residues 11 positions away from the degron boundaries. Phosphorylation sites within the degron were also counted.
- Number of degron-associated ubiquitinated lysines (UB_SITES). Degrons flanking sequences were scanned searching for ubiquitinated lysines (see below). The count represented the number of ubiquitinated lysines within 3 to 20 residues away from the degron boundaries.

In summary, we calculated eleven characteristic features for any motif instance of interest.

### Post Translational Modifications dataset

Phosphorylation and ubiquitination sites were downloaded from PhosphositePlus (Hornbeck et al., 2015) (https://www.phosphosite.org/; 04/10/2018). Non-human events were discarded.

### Random motifs of similar degron lengths

For each of the 150 annotated degron instances we randomly selected 1,000 motifs of equal aminoacid length from the human proteome. Biochemical properties of the 150,000 random motifs were next calculated. These random motifs were used as background for the calculation of the z-score of biochemical properties of annotated degron instances.

Z-score of biochemical features

For each annotated degron, we first calculated the eleven biochemical features (see Calculation of biochemical features). Next, for all annotated instances the eleven features were z-scored using the average and the standard deviation of the random motifs (see above for more information about how these motifs are generated). Random motifs whose calculation of features failed were not considered. Finally, the global distribution of degrons was obtained by averaging each feature z-score over all annotated instances.

## Degron Random Forest Classifier

### Training set

We developed a classifier to predict how much a degron motif instance resembles the biochemical features of annotated degrons. Therefore, 150 annotated degron instances encompassed the positive training set. Additionally, a set of 150 motifs was randomly selected from the simulated motifs (see Random motifs of similar degron lengths) to define the true negative set. Two different true negative sets were created. The first, named proteome-wide dataset, considers any human protein to select the 150 random motifs and does not limit the search to motifs proteins with annotated degron instances. The second dataset, named protein-specific dataset, restricts the selection of random motifs to human proteins with annotated instances (i.e., for each annotated instance similar number of random motifs from the same protein are randomly selected). In both datasets, the training is composed by 300 degron instances (i.e., 150 true positives and 150 true negatives).

### Model fitting

The raw scores of the eleven features were used as variables in the model (see Calculation of biochemical features). Two Random Forest Classifiers (RF1 and RF2) were trained using the two aforementioned datasets (n_estimators=10,000, other parameters are set to default values). Scikit-learn package (Pedregosa et al., 2011) of python was used to implement the model.

### Model evaluation

The performance of both models was measured by performing 10 independent 5-fold Stratified Cross-Validation. Each of the 10 iterations considered random splits of the training sets. The mean and standard deviation of the Area Under the Curve (AUC) across all iterations represents the performance of the classifiers. Scikit-learn package of python was used to evaluate the model.

### Feature importance calculation

Feature importance was calculated using the *feature_importances* function (with default parameters) of scikit-learn package. The variance represents inter-trees variability.

### Proteome-wide identification of instances

We predicted regular expression (regex) matches for the 29 degron motifs across 32,022 human protein sequences (see Human proteome sequences). The search resulted in 84,386 regex matches of 27 motifs across 24,994 isoforms of 15,677 genes.

### Degron-likeness of degron instances

The RFC classifier trained with the proteome-wide dataset was applied to predict the degron-likeness of the 84,386 motif instances. Each predicted instance was scored with the RFC degron probability. Differences in motif specificity might result in overrepresentation of classes with higher number of instances, penalizing those with more restrictive motifs. To mitigate this issue, we defined a motif-specific high-confidence threshold as the minimum classifier score for the annotated instances of that particular motif. For instance, on the one hand DEG_SPOP_SBC_1 had 14,732 motif matches due to its lowly specific regular expression ([AVP].[ST][ST][ST]), including 7,618 (51%) having a probability of degron greater than 0.5. After applying the motif-specific threshold (i.e., DEG_SPOP_SBC_1 = 0.84 of classifier score), there were 1,310 (8%) motifs that greatly resembled biochemical features from annotated degrons. On the other hand, DEG_Kelch_KLHL3_1 (E.EE.E[AV]DQH) had only 6 motif matches, all of them having score greater than motif-specific threshold for this motif (DEG_Kelch_KLHL3_1= 0.82).

### Annotation of predicted degron instances

The 84,386 instances were subsequently classified into four classes according to the level of evidence: motif match, novel instance, high-confidence degron instance and high-confidence degron instance with E3 interaction. The first class includes those instances with a degron probability below or equal to 0.5. The second class, novel instance, represents degron instances with a degron probability greater than 0.5 but lower than the class specific threshold. The third class, high-confidence degron instance, contains instances with a probability of degron greater than the motif-specific threshold without information about protein-protein interaction (PPi) (combined_score from STRING cutoff <300) between the E3 ligase recognizing the degron motif and the target (i.e., the sequence that matched the instance). Finally, the high-confidence degron instance with E3 interaction class, emcompasses instances with a probability of degron greater than motif-specific threshold including some information about PPi (combined_score from STRING cutoff >300). The later class represents the highest level of confidence among the predictions.

## Mapping alterations into degradation regions

### Non-synonymous mutations

We first defined a clean dataset of missense mutations from both TCGA and CCLE projects. The clean dataset included missense mutations in genes that do not harbour any other Single Nucleotide Variant (SNV) in the same sample. Applying this filtering we retrieved 1,066,092 missense mutations from TCGA and 433,336 from CCLE.

Next, each missense mutation was uniquely classified into one of the following classes: i) Mutation_Altering_Motif: mutations overlapping with a predicted or annotated degron instance. ii) Mutation_Flanking_PTM: Mutations flanking degron instances hitting residues with annotated phosphorylation sites. iii) Mutation_Flanking_UB_Lysine: Mutations flanking degron instances hitting residues with annotated ubiquitination sites. iv) Mutation_Flanking_Lysines: Mutations flanking degron instances hitting non-ubiquitinated lysine residues. v) Mutation_Flanking_Degron: Mutations flanking degron instances that do not belong to the previous classes vi) Other_Missense: any other missense mutation.

When a mutation overlaps with two different classes (e.g., a mutation that a hits a phosphorylation site next to a degron and it is itself involved in a secondary degron), the priority used to classify them was Mutation_Altering_Motif > Mutation_Flanking_PTM > Mutation_Flanking_UB_Lysine > Mutation_Flanking_Lysine > Mutation_Flanking_Degron > Other_Missense.

When a mutation overlaps with two different instances of degrons, annotated instances were prioritized over predicted ones. When the mutation overlaps with the two (or more) predicted instances they were prioritized by the highest degron probability. Finally, if a mutation overlapped with several predicted instances with similar probability of degron, the most specific instance (i.e., instance that belongs to the degron motif with lowest number of predicted instances) was chosen.

### In-frame indels

We first defined a clean dataset of in-frame indels mutations from both TCGA and CCLE. The clean dataset included in-frame insertions or deletions in genes that do not harbour any other Single Nucleotide Variant (SNV) in the same sample. Applying this filtering we retrieved 8,254 in-frame indels from TCGA and 4,580 from CCLE.

Next, each in-frame indel was classified into one of the following classes: i) In_Frame_Altering_Motif: indels overlapping with a predicted or annotated degron instance. ii) in-frame_Altering_Flanking_PTM: in-frame indels flanking degron instances hitting residues with annotated phosphorylation sites. iii) in-frame_Altering_Flanking_UB_Lysine: in-frame indels flanking degron instances hitting residues with annotated ubiquitination sites. iv) in-frame_Altering_Flanking_Lysines: in-frame indels flanking degron instances hitting non-ubiquitinated lysine residues. v) in-frame_Altering_Flanking_Degron: in-frame indels hitting residues flanking degron instances that do not belong to the previous classes vi) Other_In_Frame: any other in-frame indel mutation.

In-frame indels overlapping with multiple degron instances were prioritized following similar criteria than for the mapping of missense mutations (see above).

### 3D visualization of degrons-E3 interactions

Chimera software (Pettersen et al., 2004) was used to visualize the interaction of the N-terminal degron of NFE2L2 with KEAP1 (PDB code: 3WN7) and the BTRC degron in CTNNB1 (PDB code: 1P22). Chimera mutate_residue tool was used to generate 3D models of the NFE2L2-D29H and CTNNB1-S37C mutations.

## Statistical tests of degron alterations

### NFE2L2 KEAP1 degrons

Stability Change of missense mutations and in-frame indels overlapping with the two KEAP1 degrons in NFE2L2 including 11 flanking aminoacids was compared to both other alterations outside these regions and to the wild-type counterparts. Samples with SNV in KEAP1, high-level amplifications/deletions of NFE2L2 or alterations that disrupt the epitope of the antibody and decrease the stability were not considered. Two-sided Mann-Whitney test was used to evaluate the statistical significance.

### CTNNB1 βTRCP degron

Stability Change of missense mutations and in-frame indels overlapping with βTRCP inTRCP degron in CTNNB1 including phosphorylation sites within 11 flanking aminoacids was compared to both other alterations outside these regions and to the wild-type counterparts. Samples with SNV in any annotated upstream E3 ligases of CTNNB1, high-level amplifications/deletions of CTNNB1 or alterations that disrupt the epitope of the antibody and decrease the stability were not considered. Two-sided Mann-Whitney test was used to evaluate the statistical significance.

### CCND1 FBXO31 degron

Stability Change of missense mutations and in-frame indels overlapping with FBXO31 degron in CCND1 including phosphorylation sites within 11 flanking aminoacids was compared to both other alterations outside these regions and to the wild-type counterparts. Samples with SNV in any annotated upstream E3 ligases of CCND1, high-level amplifications/deletions of CCND1 or alterations that disrupt the epitope of the antibody and decrease the stability were not considered. Two-sided Mann-Whitney test was used to evaluate the statistical significance.

### Mutations in novel degron instances

Stability Change of missense mutations and in-frame indels overlapping novel degron instances (i.e., degron probability > 0.5) including phosphorylation sites within 11 flanking aminoacids was compared to both alterations in the same proteins but outside these regions and to the wild-type counterparts. Samples with SNV in any annotated upstream E3 ligase of the target, high-level amplifications/deletions of the target or alterations that disrupt the epitope of the antibody and decrease the stability were not considered. Due to the extreme effect of TP53 alterations, mutations in TP53 were filtered out of the analysis. Two-sided Mann-Whitney test was used to evaluate the statistical significance.

### Predicted degrons by level of evidence

Missense mutations and in-frame indels overlapping with any degron instance were stratified into three groups corresponding to its annotation level: i) alterations in annotated instances ii) alterations in novel degron instances (i.e., degron probability > 0.5) and iii) alterations in motif match instances (i.e., degron probability < 0.5). Alterations hitting phosphorylation sites within 11 flanking aminoacids were also considered. Pairwise Stability Change comparison the three groups to both alterations in the same proteins but outside these regions and to the wild-type counterparts was performed. Samples with SNV in any annotated upstream E3 ligase of the target, high-level amplifications/deletions of the target or alterations that disrupt the epitope of the antibody and decrease the stability were not considered. Due to the extreme effect of TP53 alterations, mutations in TP53 were filtered out of the analysis. Two-sided Mann-Whitney test was used to evaluate the statistical significance.

### Analysis by tumorigenic mechanism

To classify each protein according to its tumorigenic mode of action (MoA) we used the Cancer Genome Interpreter (https://www.cancergenomeinterpreter.org/genes; 20/10/2018) MoA classification. We then compared the effect of overlapping alterations in novel instances (including mutations altering the motif and mutations in flanking phosphorylation sites) in oncogenes to other alterations and and the wild-type form of the same oncogenes.

Similarly, we compared the effect of novel degron instances alterations in Tumor Suppressor Genes (TSGs) to other alterations and and the wild-type form of the same TSGs. Samples with SNV in any annotated upstream E3 ligase of the target, high-level amplifications/deletions of the target or alterations that disrupt the epitope of the antibody and decrease the stability were not considered. Due to the extreme effect of TP53 alterations, mutations in TP53 were filtered out of the analysis. Two-sided Mann-Whitney test was used to evaluate the statistical significance.

### Analysis in the MS dataset

Stability Change of missense mutations overlapping novel degron instances (i.e., degron probability > 0.5) was compared to both alterations in the same proteins but outside these regions and to the wild-type counterparts in the MS dataset. Samples with SNV in any annotated upstream E3 ligase of the target, high-level amplifications/deletions of the target or alterations that disrupt the epitope of the antibody and decrease the stability were not considered. Due to the extreme effect of TP53 alterations, mutations in TP53 were filtered out of the analysis. Two-sided Mann-Whitney test was used to evaluate the statistical significance.

## Identification of *de novo* degrons

### Selection of highly-stabilizing alterations

We first selected missense mutations and in-frame indels from TCGA that do not overlap with a degron instance -or their flanking position- and have Stability Change measures. Alterations in samples with altered upstream E3 ligases, with high-level amplifications/deletions or with alterations that disrupt the epitope region of the antibody were filtered out. A total amount of 10844 missense mutations and 288 in frame indels were selected for further analysis.

We next classified the Stability Change of the aforementioned alterations in quartiles and selected those alterations whose Stability Change is within the upper quartile. These mutations were considered as highly-stabilizing. A total amount of 2212 missense mutations and 78 in frame indels composed the highly-stabilizing alterations dataset.

### Protein profile of degron probability

For each protein with at least one highly-stabilizing alteration we performed a search for regions with degron-like biochemical properties. To do so we followed the next steps: i) First we denote k_*i*_ to an amino acid k in the i^th^ position in the sequence of a protein of length L. For all amino acids k_i_ in k_1..L_ of a protein, we first computed the degron probability from the k_i_ amino acid to k_*i+N*_ (default N=3) residues upstream. ii) Next, for each k_*i*_ in the sequence, we averaged the degron probability of all regions that contained the *i*_*th*_ position. In other words, if N=3, we averaged the degron probability of the k_i-3…i_,k_i-2…i+1_,k_i-1…i+2_ and k_i…i+3_ regions. This enabled the creation of a residue-specific profile of the degron-likeness of all amino acids of a sequence of interest. iii) We next used the profile to find peaks that represent delimited regions with favourable biochemical properties to harbour a degron. Peaks were called using the scipy find_peaks function with the following parameters width=(3, 100), distance=5, height=0.5, prominence=0.0. Peak calling resulted in 2,662 peaks in 149 proteins. iv) We finally mapped all highly-stabilizing mutations (see above) into the degron-like regions (i.e., those resulting from the peak calling). Peaks harbouring at least two highly-stabilizing alterations were selected for further analysis.

### Z-score of degron-like regions

The mean Stability Change of alterations in degron-like regions was z-scored using the Stability Change across the remainder alterations in the protein. This z-score is intended to measure how local the stabilization effect is. Regions with low z-scores (i.e., z-score below 0.5) are likely to represent non-local stabilization effects (i.e., proteins where alterations systematically stabilize the protein such as in TP53) and are subsequently filtered out. The final dataset was composed of 31 degron-like regions in 23 proteins.

### Mapping E3 ligases to targets harbouring degron-like regions

For each protein harbouring degron-like regions we annotated which E3 ligases are known to interact with the target from curated list of E3 ligase substrate interactions.

## Positive selection analysis of degrons

### Definition of the genomic regions

We confined the search of positive selection in degrons to novel degron instances (degron probability > 0.5). For each of these motif instances, regions to be analyzed contained all residues in the degron motif extended to eleven amino acid flanking positions. For instance, βTRCP inTRCP degron of CTNNB1 spanning from the 32nd to the 37th residues is encoded as P35222-1:DEG_SCF_TRCP1_1:21:48, which corresponds to its UniProt isoform ID, degron name, aminoacid flanking start, amino acid flanking end.

All aforementioned regions were subsequently mapped to GRCh37 genomic coordinates. To perform the mapping, we first retrieved, when available, the Consensus CDS (CCDS) identifier of each UniProt isoform ID. Motifs in isoforms lacking of CCDS were not further considered. Next, for all amino acids in a degron region we used TransVar (Zhou et al., 2015) to convert from amino acid position to genomic coordinates. Motif instances mapping to genomic coordinates in sex chromosomes were discarded. Genomic coordinates for 18,238 motif instances were successfully retrieved. A similar procedure was followed to retrieve the genomic coordinates of the 31 *de novo* degrons.

## Datasets of somatic alterations

### FMDeg and SMDeg

Single Nucleotide Variants (SNV) missense mutations were gathered from the TCGA and CCLE datasets (see Data collection and preprocessing for more information). For each TCGA cancer type we removed hypermutated samples. Hypermutated samples were defined as those samples with a number of missense mutations greater than 3.5 times the interquartile range plus the 75h percentile of the distribution of missense mutations in all the samples in that cohort (or in the PAN cohort for the pan-cancer analysis). Moreover, a minimum number of 1,500 missense mutations are needed to be considered hypermutated. 99 TCGA samples were considered hypermutated. A total number of 999,974 missense mutations composed the dataset of mutations.

Similarly, the CCLE dataset resulted in 62 hypermutated cell lines using the same rationale. A total number of 378,200 missense mutations composed the CCLE dataset of mutations.

### Indels

We used in-frame indels from CCLE and TCGA (see above for more information about the processing of in-frame indels) to perform the positive selection test.

### Fusions

Pan-cancer gene fusions in the TCGA cohort was gathered from the Tumor Fusion Gene Data Portal (http://www.tumorfusions.org/; 10/07/2018). We focused on fusions that conserve the reading frame (i.e., in-frame) and with a reliable level of evidence (i.e., tier 1, tier 2 or tier 3).

### SMDeg test

We developed, SMDeg, a method to identify motif instances with significantly higher number of missense mutations that expected by chance. For each query region (see Definition of the genomic regions) within a protein in a cohort of samples, SMDeg performs the following steps:

1. It first calculates the number of overlapping missense mutations with the region (denoted as O) and total number of missense mutations (named as T) that the protein harbours. By default, regions with less than 3 overlapping mutations are not further analyzed.
2. It then performs N simulations (N=10_3_ by default) were it randomly distributes T missense mutations across the protein. The simulation rationale is similar than the one employed in OncodriveFML (Mularoni et al., 2016). Briefly, the probability of each mutation to occur is based on the mutational signatures (i.e. tri-nucleotide composition) observed in the cohort. For the analysis presented in this study we pre-computed matrices of trinucleotide probabilities of the 96 possible changes across all TCGA cohorts. Matrices are computed by counting all the observed mutations in all the tumors. The random distribution of mutation will thus consider the different probabilities that each mutation has to occur.
3. Next, for each of the N simulations the method counts the number of missense mutations overlapping with the query region.
4. Once all simulations finish, It computes the average number of overlapping missense mutations across the N simulations.
5. For each input region with enough number of mutations It performs a log-likelihood test (i.e., G-test) of the number of observed overlapping missense mutations versus the mean value of overlapping mutations in the simulations.
6. Finally for each input cohort or group of cancer cell lines the resulting p-values are then adjusted with a multiple testing correction using the Benjamini–Hochberg procedure (alpha=0.05).

### DegFI score

We have developed a score that quantifies the degree of impact of missense mutations to the destruction of a degron. The score is based on observed phenotypic changes induced by missense mutations across the proteins with available RPPA alongside prior knowledge. We first assigned a baseline score to the five classes of alterations associated to degradation motifs that corresponds to the observed phenotype in our analysis (i.e., Mutation_Altering_Motif: 5.0, Mutation_Flanking_PTM: 4.0, Mutation_Flanking_UB_Lysine: 4.0, Mutation_Flanking_Lysines: 1.0 and Mutation_Flanking_Motif: 0.0). Since it is reasonable to think that all mutations overlapping with the degron do not have the same impact on degron recognition we also included features enabling the ranking of the degree of impact for the class Mutation_Altering_Motif. More specifically, we questioned whether a mutation changed degrons pattern recognition or whether it hit a phosphorylation site within the degron. Mutations of any of these types added one -or two for mutations with both properties- to their baseline score. Additionally, we measured the degree of change of two essential biochemical properties of degrons: hydrophobicity and flexibility. We measured amino acid flexibility change upon mutation as the absolute difference between the normalized Average Flexibility Index (Bhaskaran and Ponnuswamy, 1988) of the wild-type and the mutated residue. The flexibility change ranges from 1.0 for high flexibility changes (e.g., glycine to methionine mutations) to 0.0 for negligible changes (e.g., phenylalanine to tryptophan). Similarly, we defined hydrophobicity change upon mutation as the the absolute difference between the normalized hydrophobicity index (Eisenberg et al., 1984) of the wild-type and the mutated residue. Hydrophobicity change ranges from 1.0 for dramatic changes (e.g., arginine to isoleucine) to 0.0 for negligible changes (e.g., leucine to valine). All these scores were added to the baseline score of Mutation_Altering_Motif and were subsequently normalized from 1.0 (i.e., most severe degron-affecting mutation) to 0.0 (i.e., neutral mutation).

As an example, S37C, a recurrent mutation in CTNNB1 βTRCP inTRCP degron, scored 7.78 (0.86 after normalization), which is the result of the baseline score (5), the fact that it alters the pattern (+1), hits a PTM (+1), it slightly alters hydrophobicity (+0.12) and notably alters residue flexibility (+0.66). A second example in the same degron, H36P, does not change the pattern nor hits a PTM site, but it leads to considerable changes in the biochemical landscape of the degron. Therefore, it scores (0.65 after normalization), which is the result of baseline score (5), the change in hydrophobicity (+0.13) and the change in residue flexibility (+0.80).

Finally, we also weighted the impact of mutations in flanking positions by their normalized distance to their associated degron. We calculated normalized distance scores using the following formula:

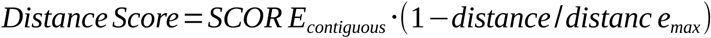

Where SCORE_contiguos_ represents the maximum score of mutations in an amino acid next to a degron (by default is equal to 3.0) and distance_max_ represents the maximum distance of flanking positions, in our case 12 residues. Normalized distance score values ranged from 0.30 to positions contiguous to the degron to 0.02 to mutations 11 residues away. The normalized distance score was added to the baseline score for each mutation.

When a mutation overlaps with two or more degrons, all possible degFI scores all calculated and the most severe score (i.e., the higher) is chosen as representative of the impact.

DegFI score was calculated for all possible missense mutations within novel degron instances (i.e., probability of degron greater than 0.5) across the genome. Mutations with high degFI scores represent mutations more likely to impair the degradation of its protein. Likewise, low degFI score represent mutations less likely to exert a change in the protein degradation.

### Validation of degFI

We computed degFI score of missense mutations overlapping with novel instances with available Stability Change. We filtered out samples with high-level amplifications/deletions, with alterations on upstream annotated E3 ligases of the gene or with alterations in the antibody epitope that decreases the stability. 388 missense mutations were successfully mapped with both degFI score and Stability Change data. We then split the mutations into two groups according to their degFI score. The first group is composed by mutations with degFI score greater than a given threshold, and the second is composed by those mutations with degFI score lower than the threshold. We used three different thresholds to evaluate the robustness of the results: the average degFI (0.21), the median degFI (0.13) and a threshold of 0.5. We then compared the Stability Change of mutations from the two groups. One-sided Mann-Whitney test was used to evaluate the significance of the difference.

### FMDeg test

OncodriveFML (Mularoni et al., 2016) is a tool that detects genes under positive selection by analysing functional impact bias in somatic mutations. Herein we defined functional impact as the effect on protein degradation measured by degFI.

OncodriveFML requieres as input the genomic regions to be analyzed. In our particular case, we used the genomic regions overlapping with degrons defined in the Definition of the genomic regions section (see above).

For each genomic region of interest with at least three missense mutations in the cohort of interest, OncodriveFML with DegFI score (hereafter named as FMDeg) simulated equal number of missense mutations within the genomic region of interest. We used 10^x8^ simulations to compute the empirical p-value. Since degFI is defined to measure the impact of single amino acid change in protein degradation other types of SNV such as synonymous mutations, splice affecting variants or nonsense mutations were not considered.

## Positive selection test for in frame indels

For each novel degron instance we evaluated whether the number of degron overlapping -- including 11 flanking residues-- in frame indels from TCGA and CCLE is statistically significant compared to the total number of in frame indels that the protein harbors in each dataset. Fisher’s exact test was used to evaluate the significance. The effect size was defined as the odds ratio (OR; see below) between the number of observed indels in the degron region (O_D_), the number of total observed indels in the protein (O_T_), the length of the degron region (L_D_) and the total length of the protein (L_T_). Degron instances with less than three in frame indels were not considered in the analysis.

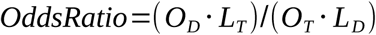

### Mapping gene fusions into degrons

We detected those gene fusions from TCGA (see above for more information about the original Fusions dataset) that lead to a total loss of a predicted instance of a degron. To do so, we used the genomic coordinates of novel degron instances used in the positive selection test. More specifically, we followed the next steps :

1. For each in-frame fusion involving two genes (i.e., gene A and gene B) we first defined the relative position of the genes in the fusion. On the one hand, Gene A is defined as the gene that conserves the N-terminus and that loses a C-terminal part of the protein. Gene fusions where the junction happens before the protein start codon of Gene A are not considered (i.e., gene fusions where there is no coding region of Gene A included). On the other hand, Gene B, is defined as the gene that conserves the C-terminal part of the protein. Gene fusions where the junction happened after the stop codon in Gene B are not considered (i.e., gene fusions that do not include any coding region of the Gene B). One gene can be labeled as Gene B in a fusion and otherwise in a different gene fusion.
2. Then for each degron instance in A genes, we annotated all fusions where the Junction in A happens before the 5’ coordinate of the first aminoacid involved in the degron region (or afterwards in the reverse strand).
3. Similarly, for each degron in B genes, we annotated all fusions where the junction in B happens after the 3’ coordinate of the last aminoacid involved in the degron region (or before in the reverse strand).
4. Finally, we counted the number of gene fusions overlapping with a particular degron instance in all TCGA samples.

## Analysis of E3 ligases alterations and the role in tumorigenesis

### Positive selection in E3 ligases

To detect E3 ligases with significant signals of positive selection in human tumor samples and cancer cell lines, we ran OncodriveFML (Mularoni et al., 2016) (default coding parameters, including indels, sampling of 10^6^ iterations and CADD (Kircher et al., 2014) 1.0) and dNdScv (with default parameters). On the one hand, OncodriveFML aims to detect genes under positive selection by analysing functional impact bias in somatic mutations. On the other hand, dNdScv detects genes under positive selection by analysing frequency of mutations compared to the background mutation rate.

To run both methods we removed hypermutated samples from TCGA and CCLE, as described above for SMDeg and FMDeg.

We filtered the output of both methods to narrow down the search to E3 ligases using the curated list of proteins involved in ubiquitination. We used a False Discovery Rate (FDR) of 10% to define as significant the signal of an E3 ligase in a particular cohort. The union of both outputs composes the set of significant E3 ligases in a dataset.

### Stability Change upon mutation in E3 ligases

We analyzed the downstream effect of nonsynonymous variants in E3 ligases to their associated degradation targets. To perform the analysis we followed the next steps:

1. We first defined the set of interacting E3 ligases-substrate pairs as the union of the annotated interactions from the curated list of E3 ligase-substrate interactions (see above) and the set predicted interactions resulting from the prediction of degron motifs across the proteome. Therefore, for each novel degron instance we included the interaction of the E3 ligase recognizing the motif and the protein matching the motif.
2. We then performed a cohort-specific analysis for TCGA tumor samples alongside a pan-cancer analysis for TCGA and for cancer cell lines.
3. For each cohort or group of cancer cell lines defined in 2) we selected those E3 ligases with at least N samples with alterations in the particular cohort (N=10 for TCGA cohorts; N=20 por TCGA pan-cancer and cell lines).
4. For each E3 ligase selected in 2), we interrogated those substrates from 1) with, at least, N wild-type samples with available RPPA data (N=10 for TCGA cohorts; N=20 por TCGA pan-cancer and cell lines). We defined wild-type samples as those samples that do not harbor a high-level amplification/deletion or single nucleotide variants in the substrate.
5. For each resulting pairs from 4) we tested whether the samples harboring mutations in the E3 ligase are significantly more stable than their wild_type counterparts. Two-sided Mann-Whitney test was used to evaluate the significance.
6. Finally for each input cohort or group of cancer cell lines the resulting p-values are then adjusted with a multiple testing correction using the Benjamini–Hochberg procedure (alpha=0.05).

### Driver mutations in E3 ubiquitin ligases

We performed an estimation of the number of mutations in excess (i.e., the difference between the number of mutations observed and the number of mutations predicted by a neutral selection model) in driver E3 ubiquitin ligases. To compute such estimation we resorted to the methodology laid out by the dNdScv method (Martincorena et al., 2017). Briefly, dNdScv provides a gene-specific estimation of the ratio of non-synonymous to synonymous substitutions (dN/dS, hereinafter termed *ω*) that allows for correction by i) chromatin features explaining the regional variability of neutral mutation rate, ii) the consequence type of the substitutions and iii) the mutational processes operative in the tumor. For our analysis we group the non-synonymous substitutions into three main consequence types: missense mutations, nonsense mutations and mutations affecting splicing.

Upon the estimation of *ω*_*c*_ for a given consequence type *c* we can estimate the number of mutations in excess for each gene-cohort and consequence type *c* as *e*_*c*_ =(*ω*_*c*_-1) *·m*_*c*_ / *ω*_*c,*_ where *m*_*c*_is the number of mutations observed with consequence type *c.* By adding the number of mutations in excess across a pool of genes (bearing signals of positive selection) we can provide an estimate of the number of driver mutations per sample. We used this rationale to calculate the number of driver mutations hitting the 28 E3 ubiquitin ligases per sample across the 33 TCGA cohorts (32 tumor types plus the pan-cancer cohort).

### Driver mutations in degrons

We performed an estimation of the number of missense mutations in excess (i.e., driver mutations) in degrons showing signals of positive selection across TCGA cohorts. We first defined a list of driver degrons, as those degrons showing significant signals of positive selection according to both FMDeg and SMDeg tests across TCGA cohorts (FDR *≤* 25%). A total of 26 genes bore degrons with significant signals of positive selection.

To estimate the number of missense mutations in excess in degrons we reproduced the analytical steps of dNdScv to compute the gene-specific *ω*_*mis*_ estimate, albeit with our own sub-genic elements of choice. Thus for any gene we were able to compute *ω*_*mis*_ for both the entire CDS and smaller coding subsequences. As degrons encompass only small stretches spanning *≤* 5 % of the total CDS length, the method lacks the sensitivity to tackle the problem by direct analysis of the degron sequence. However, we can estimate *ω*_*mis*_ for both the entire gene and the complement of the degron sequence, whence we can infer the number of missense mutations in excess for each. Finally, the number of missense mutations in excess in the degron can be given as:

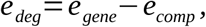

where *e*_*gene*_ and *e*_*comp*_ are the missense mutations in excess of the gene and degron complement, respectively, computed from the respective values of *ω*_*mis*_ as described in the previous section.

### Ratio of driver mutations involved in UPS

We estimated the relative contribution of driver mutations involved in UMP compared to the total driver mutations across human primary tumors from TCGA. To do so, we first performed an sample-specific estimation of the total number of driver mutations in driver genes across TCGA cohorts. We defined a dataset of 405 driver genes as the union of the 369 genes defined in the dNdScv manuscript, the 28 driver E3 ligases and the 26 driver degrons. We followed the same produce described in previous sections to perform the estimation of total of driver mutations per cohort (see above). We then performed a sample-specific ratio between driver mutations associated with UPS (i.e., either driver mutations in E3 ligases or driver mutations in degrons) and all driver mutations in that sample. This was performed across all TCGA samples analyzed.

### Stability change across all oncogenes

We performed an estimation of the stability effect of missense mutations across oncogenes with available RPPA data. We first selected oncogenes (using the mode of action classification from https://www.cancergenomeinterpreter.org/genes) with at least ten samples with missense mutations. Samples bearing mutations in annotated E3 ligases, with high-level amplification/deletions or with mutations interfering with the epitope of the antibody and decreasing the stability were not considered. For each oncogene with enough number of mutated samples we compared the Stability Change between the two group of samples (i.e., mutated and wild-type). The difference between the average Stability Change of each group of samples is used to measure the effect on the protein stability upon mutation. Two-sided Mann-Whitney test was used to measure the significance.

### Actionability of oncoproteins

We analyzed how much the actionability of oncogenes can be expanded to tumor samples bearing alterations in their E3 ubiquitin ligases. To perform this analysis we followed the next steps:

1. Sample-specific information about target actionability was extracted from the CGI. Briefly, for each tumor sample it provides information about which particular alterations are sensitive to a specific treatment (i.e, biomarkers of treatment response for that particular cancer type). We performed a cohort-specific analysis of actionability for the 32 TCGA cohorts. We used the aforementioned sets of somatic mutations, copy number alterations and gene fusions to run CGI.
2. We filtered those samples bearing biomarkers of sensitivity (i.e. EFFECT = “Responsive”) in oncogenes with available RPPA data. We used the definition of oncogenes from the CGI list (https://www.cancergenomeinterpreter.org/genes). 25 oncogenes fulfilled both conditions.
3. For each oncogene from 2) we fetched the number of samples bearing alterations in their annotated E3 ligase/s. Consequently, we defined lists of triplets <E3 ubiquitin ligase, oncogene, tumor type> where the E3 ligase had, at least, one mutation in that particular cancer type and the oncogene had been defined in, at least, one sample as biomarker of sensitivity in that cohort. 664 triplets resulted from this step.
4. For each triplet defined in 3), we defined three separate groups of samples. The first group, named as “Direct Biomarker”, is composed by samples with evidence of response. In other words, samples bearing actionable alterations in the oncogene for that particular cancer type. The second group, named as “E3 ligase biomarker with evidence” is composed by samples bearing alterations in the E3 ligase --including high-level deletions and nonsynonymous mutations-- leading to a stabilization (Stability Change greater than 0.1) of the substrate oncogene. Samples with alterations in the oncogene are filtered out of this group. Finally the third group, named as “E3 ligase biomarker without evidence” is composed by samples with E3 ligase alterations where we do not observed stabilization of the target (Stability Change lower or equal to 0.1) or we were lacking of RPPA measurements for that sample.
5. For each input cohort, plus the pan-cancer analysis, we represented the percentage of samples of each group compared to the total number of samples analyzed in that cohort.

## Acknowledgements

N.L-B. acknowledges funding from the European Research Council (consolidator grant 682398) and Spanish Ministry of Economy and Competitiveness (SAF2015-66084-R, MINECO/FEDER, UE). A.G-P. is supported by a Ramón y Cajal contract (RYC-2013-14554). IRB Barcelona is a recipient of a Severo Ochoa Centre of Excellence Award from the Spanish Ministry of Economy and Competitiveness (MINECO; Government of Spain) and is supported by CERCA (Generalitat de Catalunya). The results shown here are in whole or part based upon data generated by the TCGA Research Network.

## Author contributions

F.M.-J. prepared and carried out most analyses, including the development of their statistical framework. F.M. carried out the estimation of excess mutations and contributed to the development of the statistical framework to compute protein expression residuals. E.L.-A. contributed to the annotation of degrons and curation of antibodies of RPPA data. N.L.-B. and A.G.-P. conceived and oversaw the study. F.M.-J, N.L.-B. and A.G.-P. drafted the manuscript. All authors participated in the interpretation and discussion of results and in the final edition of the manuscript.

---

**Figure S1.**
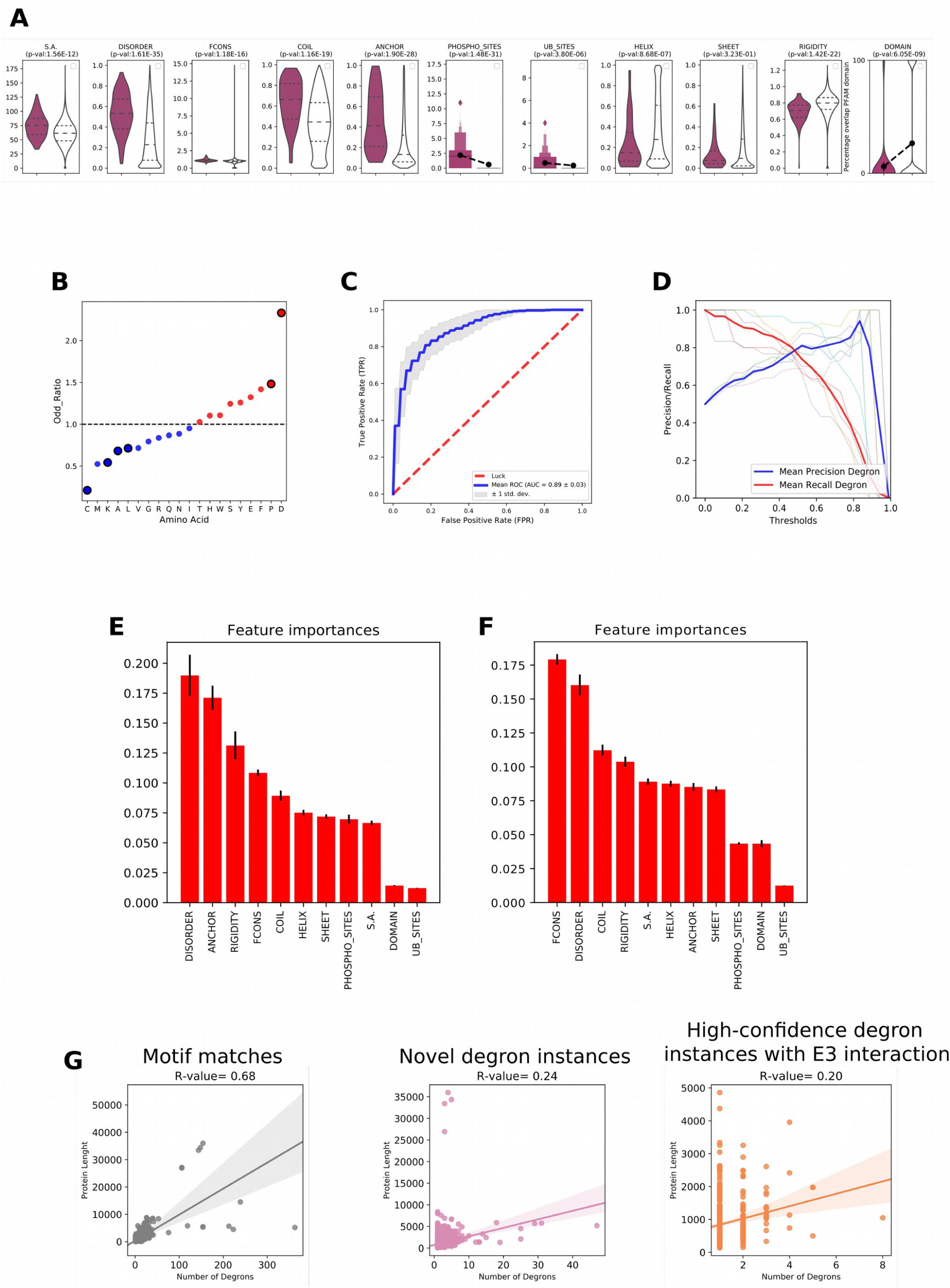
Identification of novel degron instances. (a) Distribution of the values of 11 biochemical properties of 150 annotated degron instances and the same number of aminoacid sequences of equivalent length randomly chosen from the human proteome. (b) Over or under representation of each amino acid (Fisher’s exact test odds ratio) across the sequence of 150 annotated degron instances. Significant cases (p-value<0.05) are circled in black. As described in the main paper, to identify novel degron instances we employed a random forest classifier trained on the set of 150 annotated degron instances and the same number of amino acid sequences of similar length randomly chosen from the human proteome (RF1). The ROC curve of the cross-validation of RF1 is presented in Figure 1c. To test that the results of RF1 are not due to differences in the values of the biochemical features across the sequence of proteins harboring annotated degrons and others in the proteome, we trained a second random forest (RF2) on the set of 150 annotated degron instances and the same number of amino acid sequences of similar length randomly chosen from the same set of proteins. (c) ROC curve of ten five-fold cross-validation of RF2. (d) Precision/Recall of the RF1. (e, f) Biochemical features at the top of the list of importance according to RF1 (e) and RF2 (f) (g) Correlation between the length of proteins and the number of appearances in their sequence of any motif match (left panel), novel degron instances (center panel), E3-interacting high-confidence degron instances (right panel).

**Figure S2.**
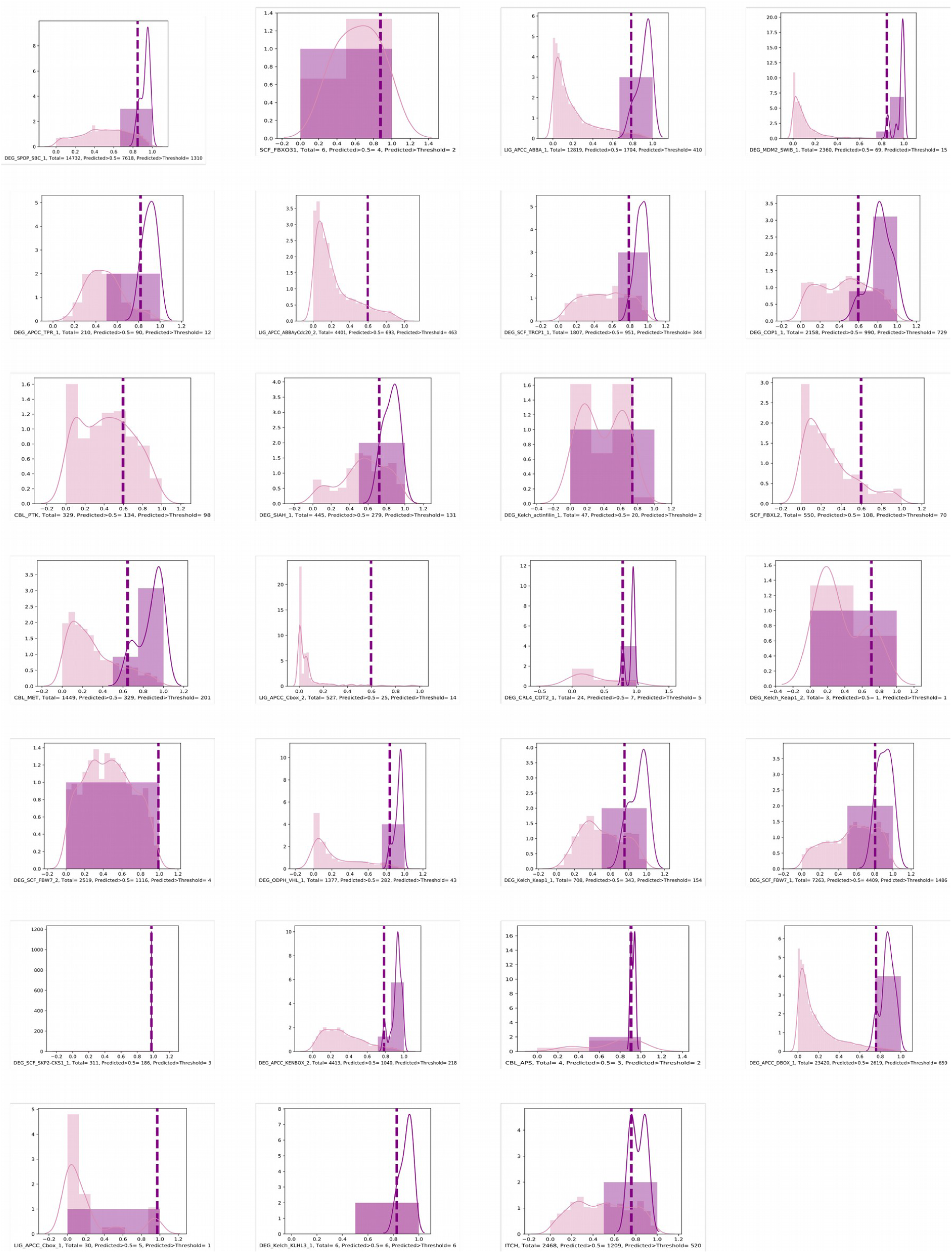
Distribution of degron probability of the matches of each motif. Each plot corresponds to the matches identified of one degron across the proteome. Degron probabilities are represented as a frequency histogram (solid light purple bars for motif matches and solid dark purple bars for annotated degron instances) and as the corresponding kernel-smoothed distribution (purple lines). Dashed vertical lines mark the site of the distribution that corresponds to the annotated degron with lowest probability, used as threshold to select high-confidence novel degron instances.

**Figure S3.**
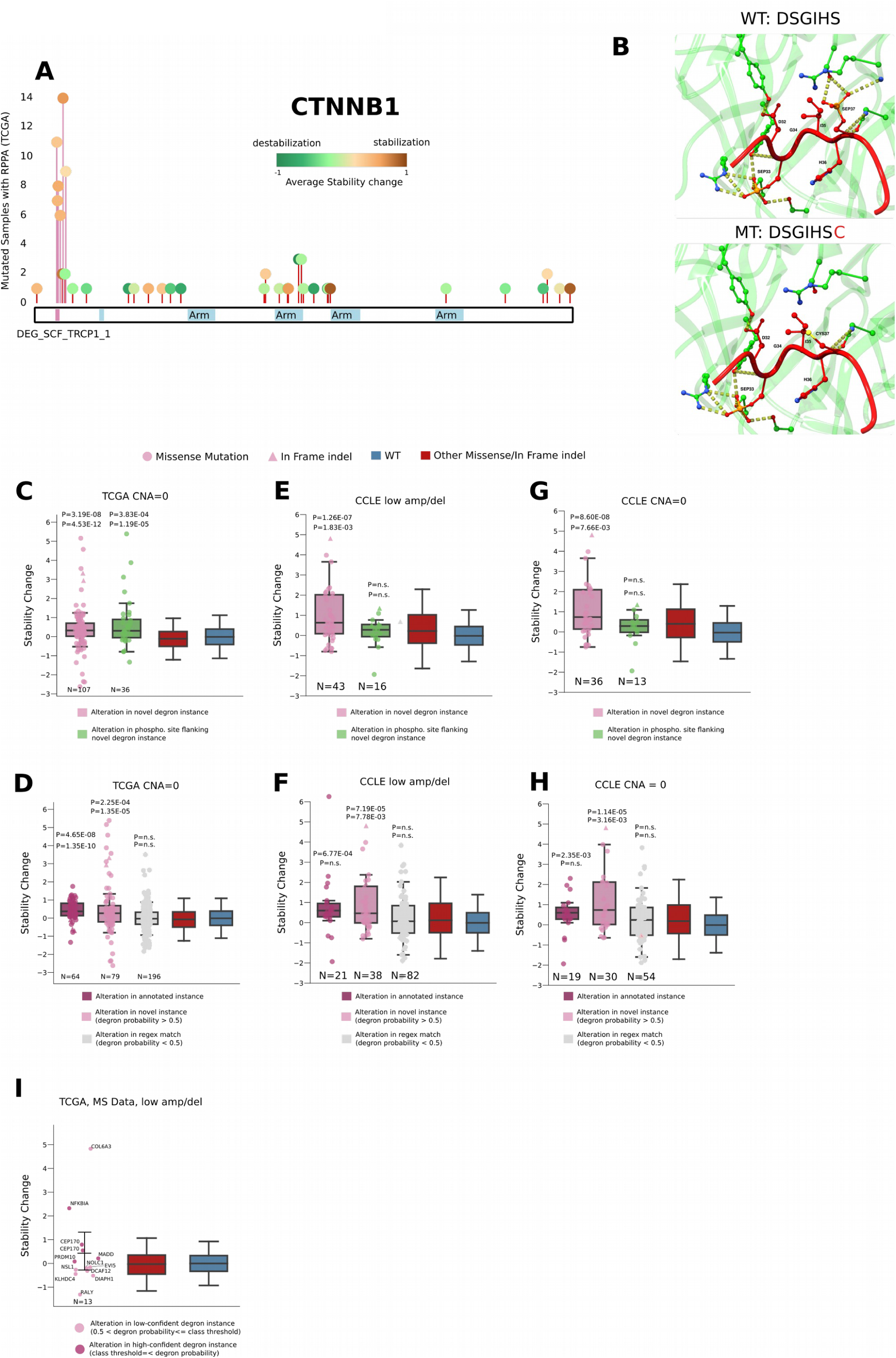
Mutations affecting degrons increase the stability of proteins. (a) Needle-plot representing the distribution of primary tumor mutations along the sequence of CTNNB1 (analogous to that of NFE2L2 in main Fig. 3a). (b) One recurrent mutation (a change of a cysteine for a serine) abrogates three hydrogen bonds in the interface of interaction between CTNNB1 and the degron of βTRCP. Chimera softwareTRCP. Chimera software (Pettersen et al., 2004)was used to visualize the CTNNB1 βTRCP. Chimera softwareTRCP interaction (PDB: 1P22). (c, d) Comparisons of protein stability upon mutations analogous to those represented in main Figures 4e and f, restricted to tumors in which the protein harboring the degron under analysis is diploid. (e, f) Comparisons of protein stability upon mutations analogous to those represented in main Figures 4e and f, but carried out using cancer cell lines mutations. (g, h) Comparisons of protein stability upon mutations analogous to those represented in panels e and f, restricted to tumors in which the protein harboring the degron under analysis is diploid. (i) Few (13) proteins carrying mutations in novel degron instances exhibit a clear trend towards stability increase (determined using mass-spectrometry rather than RPPA as in previous examples), although non-significant due to lack of statistical power.

**Figure S4.**
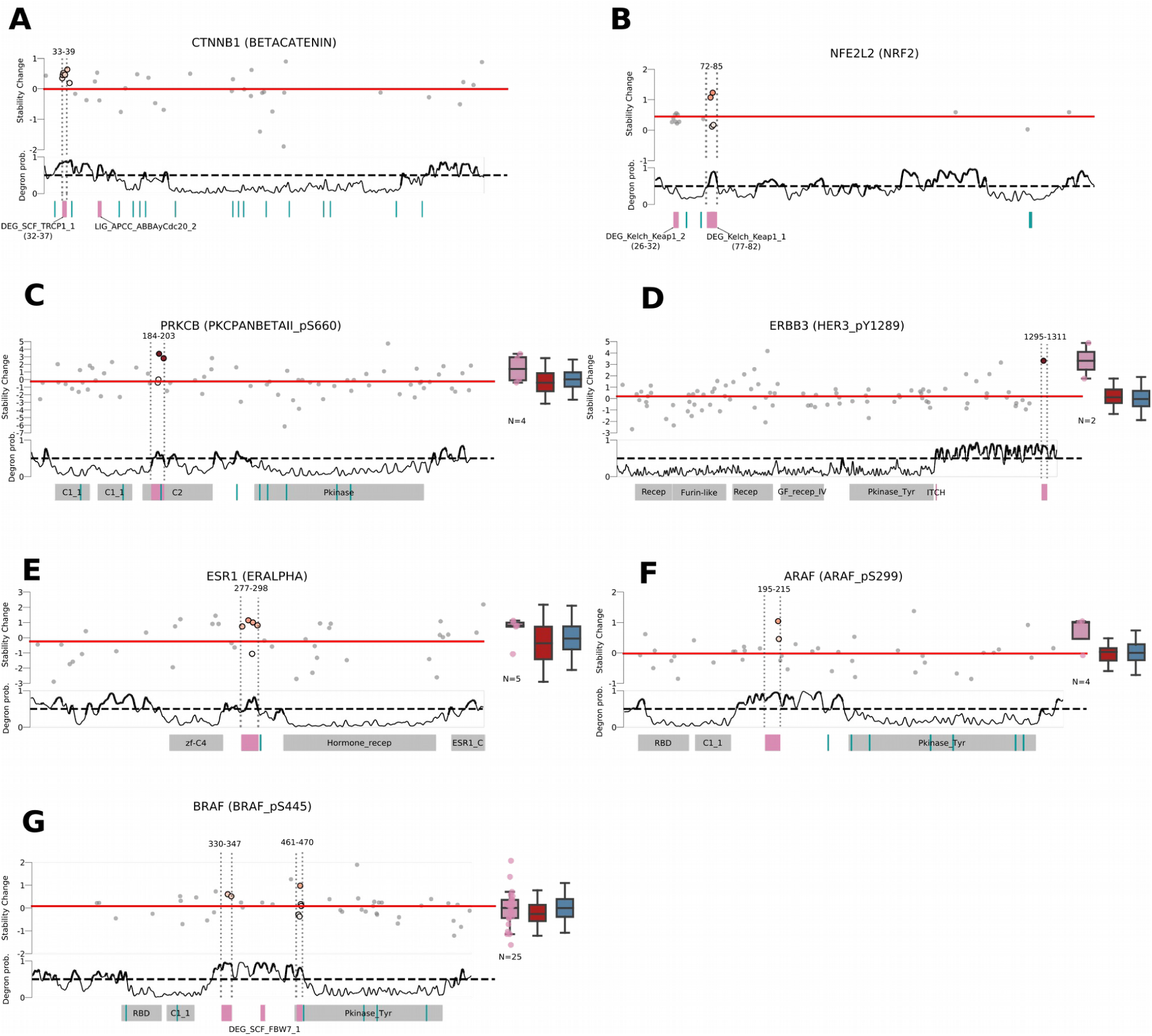
Examples of *de novo* degrons. (a,b) Identification of annotated degrons in CTNNB1 and NFE2L2 using the approach devised to identified *de novo* degrons. (c-g) Examples of *de novo* degrons identified in PRKCB (c), ERBB3 (d), ESR1 (e), ARAF (f), and BRAF (g), represented as in Figures 4c and d.

**Figure S5.**
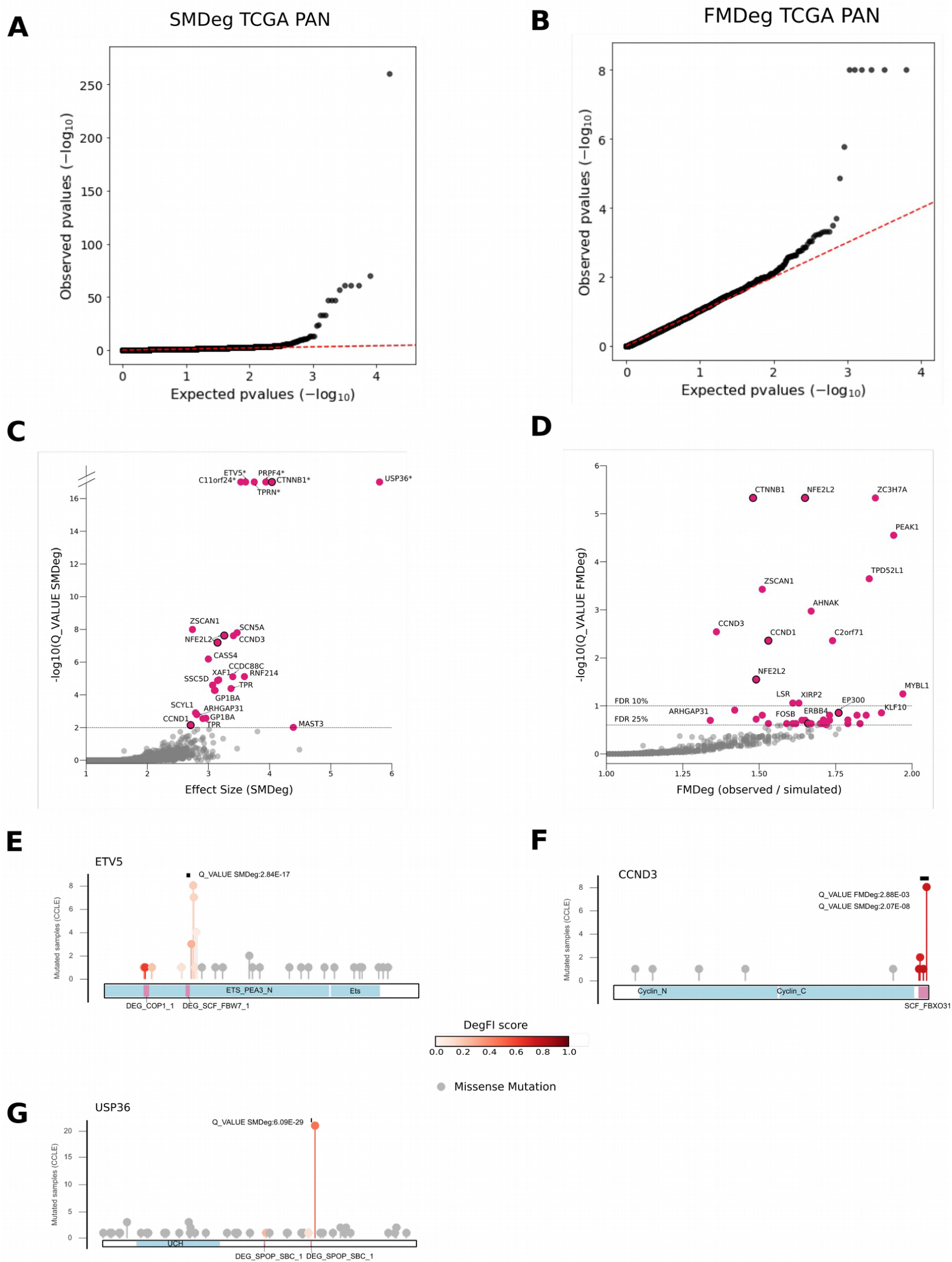
Positive selection in degrons. (a, b) QQ-plots relating the observed and expected distributions of p-values produced by the SMDeg (a) and FMDeg (b) tests on the TCGA PAN cohort. (c, d) Novel degron instances that appear significant (FDR < 1%) in the SMDeg (c), or significant (FDR < 10%) or nearly significant (FDR < 25%) in the FMDeg test (d) across cancer cell lines. (e, g) Needle-plots representing the distribution of mutations in cancer cell lines along the sequences of ETV5 (e; significant in SMDeg), CCND3 (f; significant in SMDeg and FMDeg), USP36 (g; significant in SMDeg). The representation is analogous to that presented in main Figures 5f-i.

**Figure S6.**
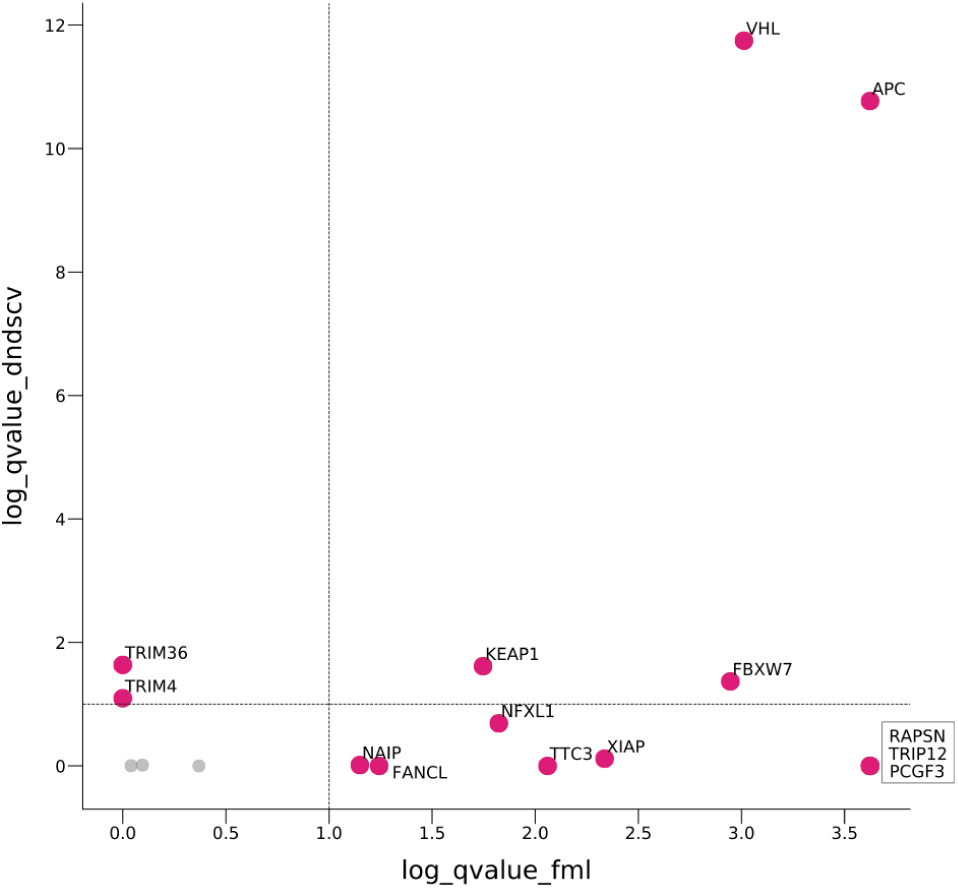
Driver E3s. Driver E3s across cancer cell lines are identified through signals of positive selection detected by OncodriveFML (Mularoni et al., 2016) and dNdScv (Martincorena et al., 2017). Analogous to main Figure 6a.

**Figure S7.**
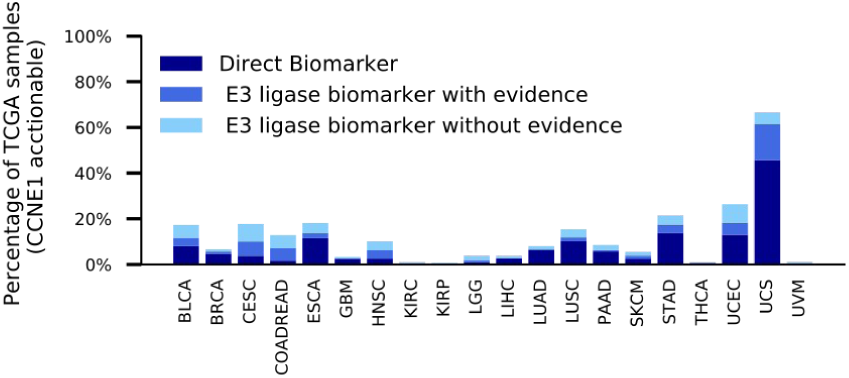
TCGA tumors with actionable UPS alterations related to CCNE1. The bars represent the proportion of tumors in each cohort with CCNE1 alterations that could be targeted directly via CDK inhibitors (dark blue), or with alterations of FBXW7 with (medium blue) or without (light blue) increased stability of CCNE1 which could in principle be targeted indirectly.

## Supplementary Note

### Protein-mRNA linear regression and residuals calculation

This supplementary note has two objectives: i) to illustrate the need for classification of CCLE antibodies into unimodally or bimodally distributed. Through the example of CDH1 we carefully explain all steps followed to perform such classification ii) to illustrate the need for normalization of raw residuals to be able to compare the Stability Change values across different proteins and/ or cohorts. We provide the explanation alongside an example to illustrate with a real example such need.

### Classification of CCLE antibodies as unimodally or bimodally distributed

Due to the inability to group samples by cohorts in the CCLE dataset, all cell lines were joined to perform the robust linear regression by IRLS. However, this grouping may result into heterogeneous distribution of mRNA-protein abundance due to tissue-specific expression patterns. More specifically, we observed that some cell lines displayed a bimodal distribution consistent with low versus high expression profiles. Figure Supplemental Note 1 illustrates this particularity thought the example of CDH1. Figure Supplemental Note 1a shows that CDH1 has a bimodal distribution across CCLE cell lines. To address this issue, each antibody was first classified into unimodally or bimodally distributed. This was done by fitting a two-dimensional Gaussian mixture model of the joint distribution of mRNA and RPPA. Antibodies whose distribution were more accurately fit by two components were classified as bimodal. The thresholds defined to classify as bimodal were Bayesian Information Criterion (BIC) difference between the models (i.e., the differences in the BIC score of the model with one component minus the bimodal model) greater than 150 and euclidean distance between centroids of the model with two distributions greater than 2.5. For antibodies classified as bimodally distributed, the Gaussian mixture model assigned to each cell line the closest distribution. Figure Supplemental Note 1b represents the classification of cell lines into two components according to the their RPPA and mRNA value for the CDH1 antibody. Finally, for both unimodals and the two independent distributions of bimodal antibodies, robust regression was performed as previously described. Figure Supplemental Note 1c represents the robust regression on the two independent distributions of CDH1 across CCLE cell lines. Samples with somatic mutations, high-level amplifications or alterations in upstream E3 ligases were not considered in the classification nor the example of CDH1. Scikit-learn package of python was used to perform the classification.

**Supplementary Note Figure 1.**
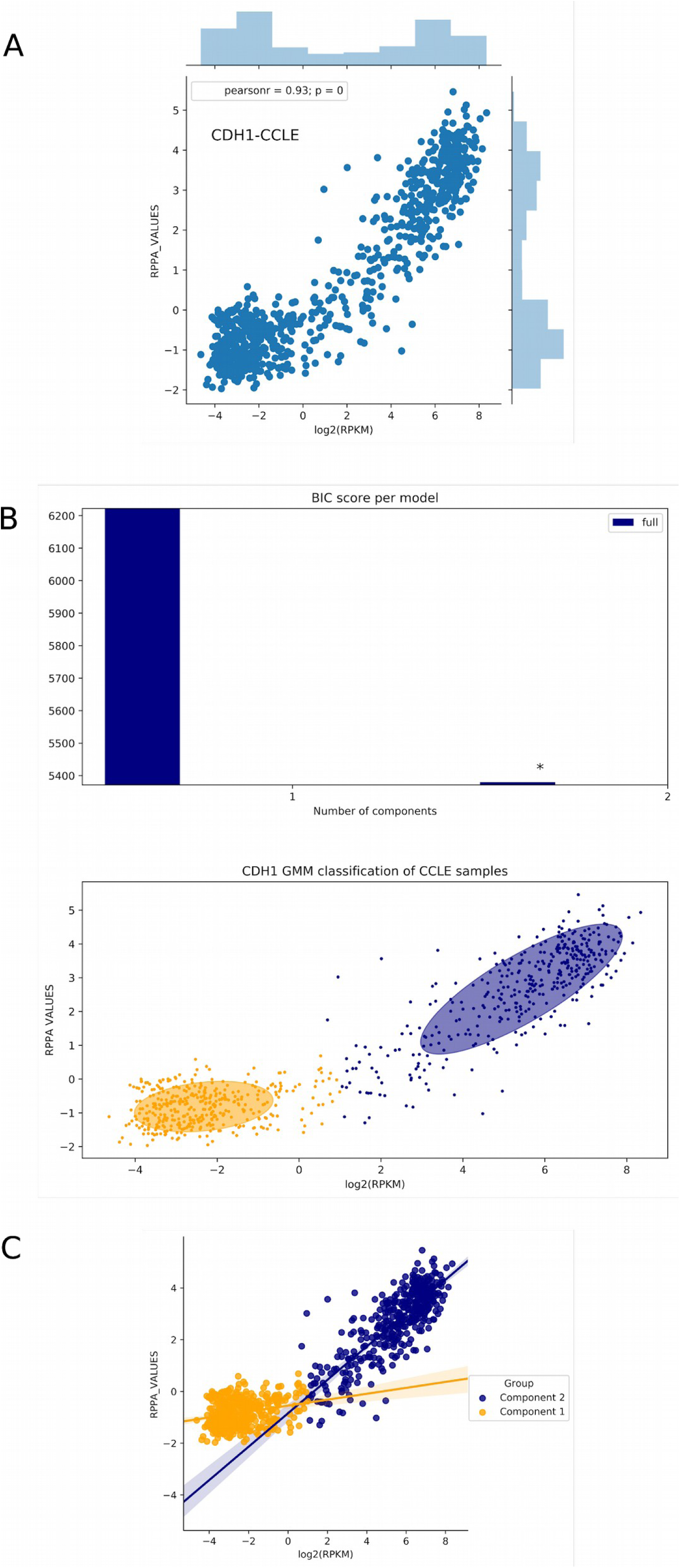
Illustrative example of CDH1 bimodal distribution across CCLE samples. a) Distribution of CDH1 mRNA expression (in log_2_ (RPKM) and protein expression (in RPPA) across CCLE samples. b) Top panel represents the BIC score for one or two components models. Bottom panel represents the two components distribution of CDH1 across CCLE. Samples orange belong to the first component while purple samples are those that belong to the second component. c) Resulting double IRLS after classification of the CDH1 distribution of bimodal. Each sample Stability Change is calculated based on its associated component IRLS.

### Raw residuals normalization across primary tumors and cancer cell lines

Raw residual were first normalized dividing each observed value by the cohort’s RPPA -or Log_Ratio(iTRAQ)-standard deviation (std). Example 1 (Supplemental Note 1a,b,c) justifies the need to apply such normalization. In this example, two pairs of protein-cancer types (EPPK1-COADREAD and CDK1-PRAD) with similar log2(RPKM) dispersion showed distinct RPPA dispersion (EPPK1 std=1.13; CDK1 std =0.04). Such differences have an influence on the distribution of raw residuals (Supplemental Note 1c, left panel). After dividing the raw residual by the RPPA std, both cases show similar distribution of residuals (Supplemental Note 1c mid and right panels).

Likewise, Example 2 (Supplemental Note Fig. 1d,e) justifies the need to normalize by mRNA expression. Lower mRNA dispersions will always tend to make higher regression slopes, thereby pushing the residuals of the mRNA outliers up. In Example 2 presents two antibody tumor type pairs (PGR in BRCA R=0.76; p-value∼0; ANXA7 in PRAD R=0.34, p-value=1.8e-10; Supplemental Note 1d). We can observe that the stds of both axes in the second pair significantly differ from the stds in the first pair. This implies that the raw residuals are accounting for this imbalance. To rule out this distortion of the raw residuals, we needed to rescale also by the mRNA std. In sum we defined a normalized residual, referred to as Stability Change:

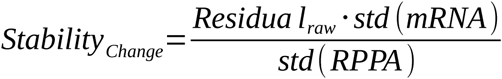

**Supplementary Note Figure 2.**
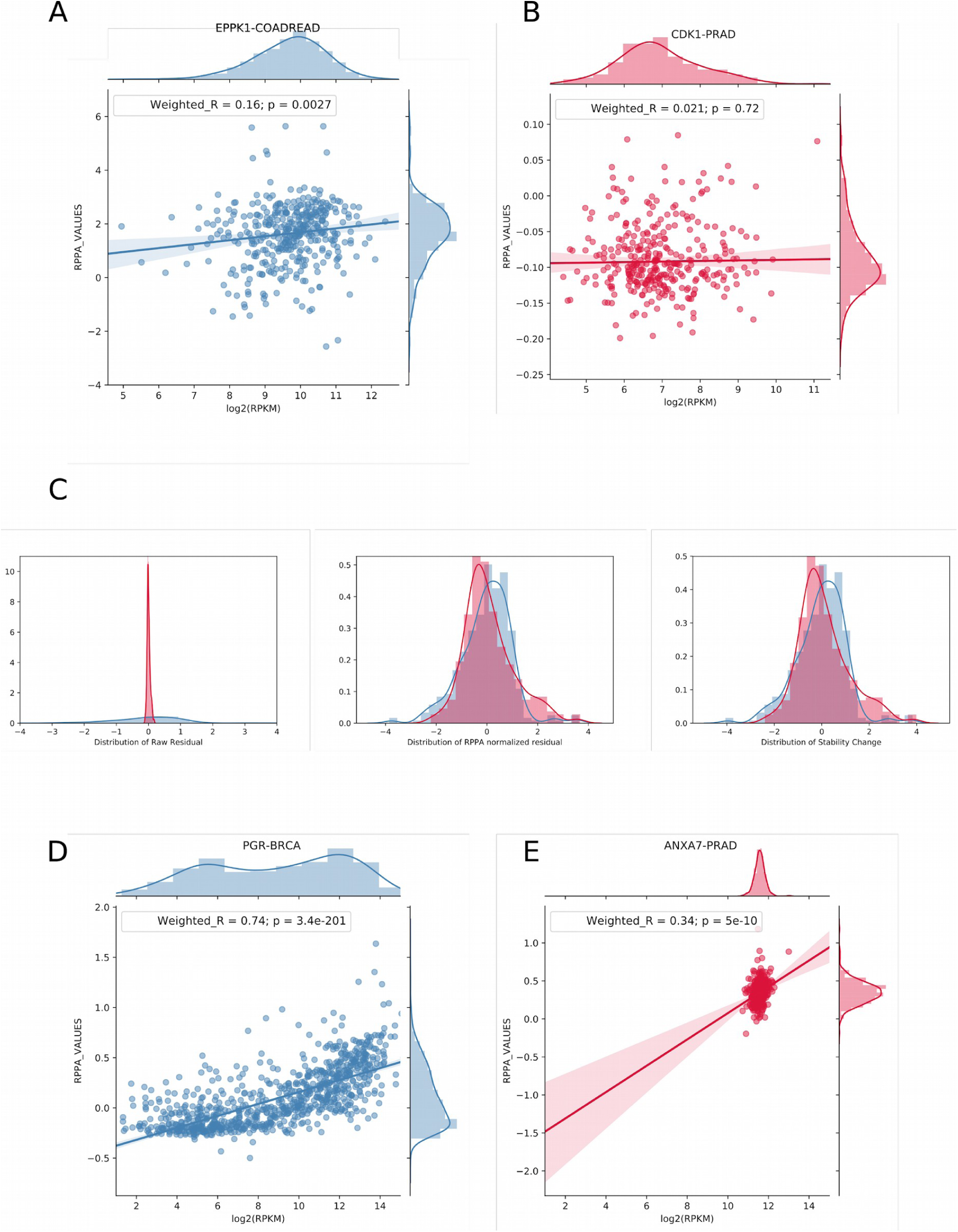
Examples that illustrate the normalization of RPPA and mRNA values across primary tumors and cancer cell lines. a) IRLS regression of epiplakin 1 (EPPK1) across colorectal adenocarcinoma (COADREAD) samples from TCGA. b) IRLS regression of CDK1 across prostate adenocarcinoma samples (PRAD) from TCGA c) Left panel shows the distribution of raw residual of EPPK1 (blue) and CDK1 (red) across the aforementioned cohorts. Mid panel shows the distribution of of raw residuals normalized by RPPA. Right panel shows the Stability Change of both distributions. d) IRLS regression of progesterone receptor (PGR) across breast cancer (BRCA) samples from TCGA. e) IRLS regression of annexin VII (ANXA7) across PRAD samples from TCGA.

